# Regulated bacterial interaction networks: A mathematical framework to describe competitive growth under inclusion of metabolite cross-feeding

**DOI:** 10.1101/2023.02.09.527847

**Authors:** Isaline Guex, Christian Mazza, Manupriyam Dubey, Maxime Batsch, Renyi Li, Jan Roelof van der Meer

**Affiliations:** Department of Mathematics, University of Fribourg, Chemin du Musée 23, CH-1700 Fribourg, Switzerland; Department of Fundamental Microbiology, University of Lausanne, CH-1015 Lausanne, Switzerland

## Abstract

When bacterial species with the same resource preferences share the same growth environment, it is commonly believed that direct competition will arise. A large variety of competition and more general ‘interaction’ models have been formulated, but what is currently lacking are models that link mono-culture growth kinetics and community growth under inclusion of emerging biological interactions, such as metabolite cross-feeding. In order to understand and mathematically describe the nature of potential cross-feeding interactions, we design experiments where two bacterial species *Pseudomonas putida* and *Pseudomonas veronii* grow in liquid medium either in mono- or as co-culture in a resource-limited environment. We measure population growth under single substrate competition or with double species-specific substrates (substrate ‘indifference’), and starting from varying cell ratios of either species. Using experimental data as input, we first consider a mean-field model of resource-based competition, which captures well the empirically observed growth rates for mono-cultures, but fails to correctly predict growth rates in co-culture mixtures, in particular for skewed starting species ratios. Based on this, we extend the model by cross-feeding interactions where the consumption of substrate by one consumer produces metabolites that in turn are resources for the other consumer, thus leading to positive feedback loops in the species system. Two different cross-feeding options were considered, which either lead to constant metabolite cross-feeding, or to a regulated form, where metabolite utilization is activated with rates according to either a threshold or a Hill function, dependent on metabolite concentration. Both mathematical proof and experimental data indicate regulated cross-feeding to be the preferred model over constant metabolite utilization, with best co-culture growth predictions in case of high Hill coefficients, close to binary (on/off) activation states. This suggests that species use the appearing metabolite concentrations only when they are becoming high enough; possibly as a consequence of their lower energetic content than the primary substrate. Metabolite sharing was particularly relevant at unbalanced starting cell ratios, causing the minority partner to proliferate more than expected from the competitive substrate because of metabolite release from the majority partner. This effect thus likely quells immediate substrate competition and may be important in natural communities with typical very skewed relative taxa abundances and slower-growing taxa. In conclusion, the regulated bacterial interaction network correctly describes species substrate growth reactions in mixtures with few kinetic parameters that can be obtained from mono-culture growth experiments.

**Author summary:** Correctly predicting growth of communities of diverse bacterial taxa remains a challenge, because of the very different growth properties of individual members and their myriads of interactions that can influence growth. Here we tried to improve and empirically validate mathematical models that combine theory of bacterial growth kinetics (i.e., Monod models) with mathematical definition of interaction parameters. We focused in particular on common cases of shared primary substrates (i.e., competition) and independent substrates (i.e., indifference) in an experimental system consisting of one fast-growing and one slower growing Pseudomonas species. Growth kinetic parameters derived from mono-culture experiments included in a Monod-type consumer-resource model explained some 75% of biomass formation of either species in co-culture, but underestimated the observed growth improvement when either of the species started as a minority compared to the other. This suggested an in important role of cross-feeding, whereby released metabolites from one of the partners is utilized by the other. Inclusion of cross-feeding feedback in the two-species Monod growth model largely explained empirical data at all species-starting ratios, in particular when cross-feeding is activated in almost binary manner as a function of metabolite concentration. Our results also indicate the importance of cross-feeding for minority taxa, which can explain their survival despite being poorly competitive.

## 2 Introduction

Bacteria and other microorganisms colonize practically any habitat on our planet, be it host-associated or in free-living environments [1]. Typically, they occupy their habitats as multi-species communities, that develop as a consequence of their seeding history and migration priorities (i.e., which species entered into the habitat at which moment) [2–4], and under the physico-chemical conditions prevailing in the habitat [5–7]. Multi-species communities intrinsically develop interspecific interactions [8–10], arising from the spatial constellations of cells from different taxa [11, 12], their physiological and metabolic properties [13,14], and distance- or contact-dependent biological mechanisms [15,16]. The large physiological and metabolic flexibility and genotype richness make it difficult to determine the main factors underlying community development, and consequently, to capture those factors in appropriate predictive mechanistic models.

A large variety of community models has been put forward over past years, which emphasize either population development within communities from Lotka-Volterra-type interactions, or are based on McArthur consumer-resource theory [17]. Models can be deterministic and population-based [18–20], or cell-oriented [21, 22] and including spatial or flux components [23–25]. Of major importance for the Lotka-Volterra models is the estimation of pair-wise interaction coefficients, the true nature or ‘strength’ of which is mostly not known, but can be estimated from model fitting. For example, field observations of plant or animal species distributions have been used to infer spatial correlations [26]. In addition, species interactions have been modeled from interacting patches [27–29] or scale transitions [26,30–32]. In case of microbial communities, interactions are frequently inferred indirectly from longitudinal measurements of relative taxa abundances within community samples, or are parameterised from comparison of single versus paired taxa growth [19,22,33,34]. The general types of interactions occurring between community members can be described from their (positive, negative or neutral) signs at steady state [35]. Lotka-Volterra models typically do not take into account substrate utilization, which governs microbial cell physiology and growth, nor the effects of the environmental or host boundary conditions. Furthermore, they rarely include system feedbacks or multi-species interactions.

In contrast to Lotka-Volterra models, which mostly function as a ‘top-down’ approach on experimental or field observations, agent-based models consider microbial cells as mathematical objects (with or without physical size), whose division and population growth is controlled by local substrate concentrations and physico-chemical conditions [23–25]. The spatial environment is broken down into grids, for which the local substrate and conditions can be calculated, and onto which the agents are placed. Agents are given metabolism and cell division properties (i.e., Monod kinetics) [36], or other biological properties such as motility or chemotaxis (i.e., Patlak-Keller-Segel model) [37]. Depending on local conditions, the agents will deplete substrate, divide and change the grid concentrations, and will occupy different positions on the surface. Concentrations and agent positions are calculated for every time step, from which the species growth or movement on the surface can be quantified. Species interactions can be introduced by defining interaction parameters that influence Monod kinetics on the substrate [36]. Agents can be given more complex metabolic functioning by including individual reduced genome-scale metabolic models [38]. This enables modeling of substrate uptake, and efflux and exchange of metabolites between species. Additional complexity, for example, from cell crowding, can be included to more realistically represent multicellular structures on surfaces. The individual based models have been further adapted to 3D porous environments, to include species distributions in space [39].

Developing a good mathematical model to describe bacterial species interactions is crucial for community growth predictions, but depends on the level of intended detail and granularity in the system (e.g., system- or agent-based), the specific questions being asked and the types of available data. Particularly for microbe-centered community models it would be important to connect to classical theory of microbial growth kinetics and physiology, such that empirical measurements of the latter (e.g., growth rate, yields) can more easily be included, but such models are currently lacking. Furthermore, there is no consensus yet on how to mathematically best link growth kinetic models of pure cultures and that of multi-species mixtures, under inclusion of emerging biological and metabolic interactions [17]. The primary objective of the work presented here was thus to describe a mathematical basis for resource-limited community growth under influence of interspecific interactions. On the basis of forward regulatory feedback theory, we propose a generalized interaction parameter, which connects to the individual species’ maximum specific growth rate. Our basic hypothesis is that population growth rates within bacterial communities are mostly influenced by carbon and nutrient availability (and less so by biological warfare mechanisms between species), and thus strongly dependent on consumption of primary substrate(s) in the system, as well as on production and utilization of metabolites (byproducts) from the primary substrates (see, for example, Ref. [40]).

We develop the model notions for the case of a two-species system, assuming that once such interaction terms are defined and parameterized, it will be easier to extend their description and usage to higher order mixtures and conditions (e.g., as in [40]). Our general population growth model is based on utilisation of a single growth-limiting resource in batch culture, which we extend by including metabolic interactions using a chemical reaction network (CRN) [20], which differs from so-called generalized consumer-resource models ([40], [17]). We then introduce metabolite-based cross-feeding where the consumption of a substrate by one of the consumers produces metabolites that can act as resources for the other, leading to positive feedback in the two-species system. We prove mathematically that such positive feedback loops cannot be constant since they would lead to vanishing steady state metabolite concentrations. This is in contradiction to previous data (see, e.g., [40] where steady-state waste concentrations were shown to persist) and results in violation of thermodynamic principles. Instead, we introduce the concept of activation thresholds, in analogy to previous gene regulatory network models [41–43], to make the positive feedback loops conditionally dependent on (summed) metabolite concentrations. Model predictions were tested and compared to experimental resource-limited batch growth with two species, one of which a fast (*Pseudomonas putida*) and the other a slower-growing bacterium (*Pseudomonas veronii*). Both species were grown individually or in co-culture in chemically defined medium, using different starting cell ratios and abundances. Population growth rates were inferred from exponential phase, and overall biomass yields from stationary phase measurements. We further experimentally impose two types of substrate scenarios, either using a primary single competitive resource (i.e., succinate) for both bacterial species, or invoking substrate ‘indifference’, in form of two substrates specific for each of the two bacteria. Data and model simulations indicate that utilization of excreted metabolites is a non-negligible part of competitive interactions. Contrary to intuition, this leads to minority populations proliferating better than expected from growth kinetic differences in competition. This effect may thus help to understand why slow-growing taxa in natural communities are persisting even under substrate competition.

## 3 Results

### 3.1 Development of a regulated community growth interaction model

#### 3.1.1 Resource competition without cross-feeding

We use the notions and tools from chemical reaction network theory (CRN; see Materials and Methods) to expand community growth kinetic models with interspecific interactions that assume metabolic cross-feeding. The basic growth kinetic parts for the population growth of each species follow logistic or Monod growth, as we will explain in brief.

The logistic model assumes that growth can be described as a reaction involving a (cell of a) species *S* that enters in contact with a resource *R*, which it transforms to new cell and leading to cell division (duplication). This process is described in general terms

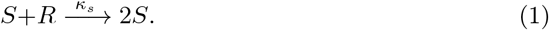

with *κ_s_* representing the rate constant. During cellular metabolism and biosynthesis, substances other than cell biomass are produced [44–48], which can leak outside the cell and which we (collectively) denote by *W*. Leaking substances may consist of regular metabolites in temporary overflow (which the cell may take up again at a later stage), or specifically excreted compounds with a biological distinct function (e.g., signalling molecules or toxins), or wastes (i.e., compounds not further used by the species). In general terms, the waste formation reaction would then look like:

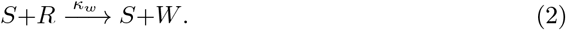

with *κ_w_* representing the respective rate constant. The CRN described by (1,2) leads to a classical **logistic model**, see Appendix S1.

In contrast to the logistic model, the **Monod model** assumes that a bacterial species *S* uses a unique resource *R* to form an intermediate complex *P*, which then gives rise both to a new cell and to waste product *W*. In this case, the CRN is reformulated as

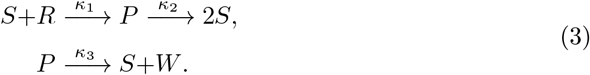

*κ*_1_, *κ*_2_, and *κ*_3_ in equation (3) refer to reaction rates, but can be deduced from the typical measured growth kinetic parameters during batch growth (i.e., yield, *μ_max_* and *K_S_*, see Appendix S2). The related o.d.e. is given in Appendix S2.

These CRNs can be extended to a community of two bacterial species *S*_1_ and *S*_2_, the related complexes *P*_1_ and *P*_2_, and a single common resource (R) with species-specific wastes *W*_1_ and *W*_2_. We show here the Monod version - see Appendix S1 for the logistic case. Further Monod model details are presented in Appendix S2.

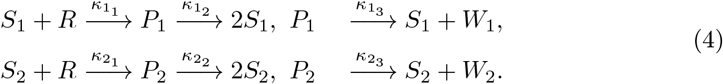

As we show below in Section 3.2.1, the Monod model (3) captures correctly the empirically observed growth rates in mono-culture (see Figure 2) but fails to explain experimental growth data in mixtures in case of imbalanced starting species ratios, see Figure 4C and Figure 6A,B.

#### 3.1.2 Resource indifference in co-cultures

We can extend the two-species resource-competition based model for co-culture growth with exclusive substrates for each of the species *R*_1_ and *R*_2_. In this case, the CRN takes the form

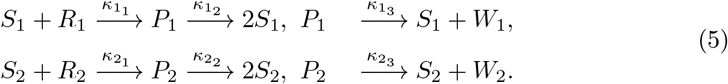

As in this case there is no competition for resource, the mass action kinetics would lead to biomass growth of species *S*_1_ and *S*_2_ with the same steady state values in co-culture as in mono-culture. We show in Section 3.3 that such mathematical predictions are valid experimentally for co-cultures with dual exclusive substrates.

#### 3.1.3 Resource competition with cross-feeding

Until here, we assumed that both species only grow on the primary resource R and produce species-specific wastes *W*_1_ and *W*_2_, which are not re-used. We can expand the models to allow utilization of the waste, in other words: to allow cross-feeding to occur. Such chemical interactions have been considered in [17,40] for generalized consumer-resource models and also in [20] in relation to Lotka-Volterra pairwise interaction models. Notably, species-specific wastes may also be re-utilized by the species in question (for example, in case of temporarily excreted metabolic intermediates, see, e.g., Ref [44]), but this is implicit in yield measurements of mono-culture growth and is therefore not considered here. We model the cross-feeding by adding the following reactions to the CRN (4)

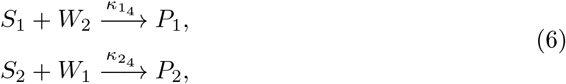

which induces two positive feedback loops. This model can be simplified by eliminating the intermediate species *P_i_, i* = 1, 2, by assuming a quasi steady state approximation. The reader can consult [41] where a full description of such standard simplification is provided. The resulting CRN is then given by

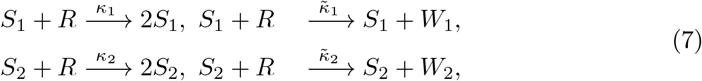

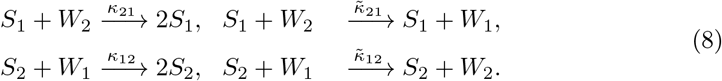

The cross-feeding CRNs (7,8) can be represented graphically as a mixture of Lotka-Volterra (LV) predator-prey interactions (see Figure 1A), corresponding to the reactions of the left-hand side in (7,8), and of catalytic conversion reactions (Figure 1B), which are given in the right-hand side of (7,8). Under consideration of mass action kinetics, this CRN provides an example of a cross-feeding mechanism where the consumption of an item of some resource (here the species *R,W*_1_ or *W*_2_) by consumers *S*_1_ or *S*_2_ produces an item which is a resource for the other consumer. However, allowing permanent cross-feeding as in the CRNs (7,8)will lead to vanishing steady-state metabolite (waste) concentrations, as shown in Propositions 1 and 2 of Supplementary Appendix S3, which is in contradiction to experimental results from literature (steady-state waste concentrations do not vanish, see, e.g. [40]). Therefore, we introduce a concentration-dependent ‘threshold’ on waste utilization, as described below.

**Figure 1.**
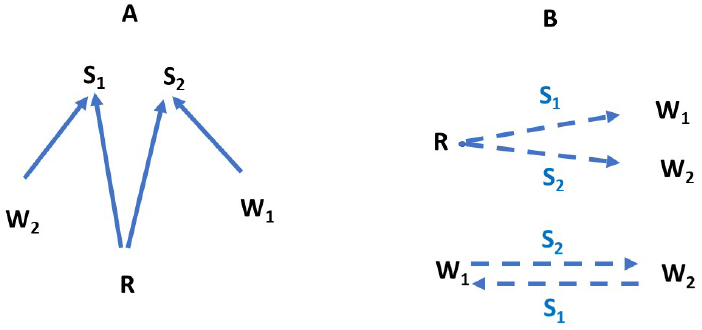
(A) The predator-prey Lotka-Volterra subsystem where species *S*_1_ preys on *R* and *W*_2_ and *S*_2_ preys on *R* and *W*_1_. (B) The catalytic conversion subsystem involving species *R, W*_1_ and *W*_2_. The species associated to the dashed arrows indicate the nature of the catalyst, and the direction of the arrow indicates the transformation product.

#### 3.1.4 Regulated bacterial cross-feeding network (RBN)

Metabolites are constantly produced by growing cells and some of those may be excreted to the outside as a result of leakage, balancing energy overflow or improperly expressed pathways [44–48]. However, depending on the growth environment and the concentration of producer cells, such metabolite concentrations may be too low to be taken up by the cross-feeding species [36]. In particular, at the onset of batch growth with low starting cell numbers, high substrate concentrations and a large medium volume, the appearing metabolite (waste byproducts) concentrations will be immediately diluted by molecular diffusion (see the experimental scenarios presented below). To account for the fact that metabolite concentrations are too low for efficient uptake by cross-feeding cells, and have less energy content than the primary substrate, we introduce a concentration-dependent threshold function for the utilization of the waste. At the start of growth, both species preferentially only consume the primary resource because of its higher concentration and it will take some time for the waste to appear in the system before being detected and used by the (respective) other species.

We can take this effect into account by constraining the utilization of waste as growth resource by a threshold function (with thresholds *T*_*W*_2__ and *T*_*W*_1__). In the case of the two-species community from above, this would lead to a modification of equation (6)

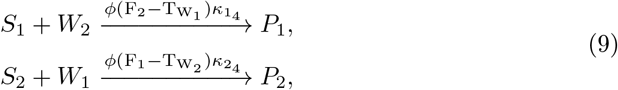

with an activation function *ϕ*(*x*) with *ϕ*(*x*)≈0 when *x* < 0 and *ϕ*(*x*)≈1 when *x* > 0, which is dependent on the differences between the concentrations *F_i_* of wastes *W_i_* and their thresholds *T_W_i__*. In the following, we develop and simulate two types of threshold functions. In the first case, the threshold function is discontinuous, whereas in the second, waste is utilized as a continuous function of its concentration. In the first case (discontinuous threshold) we take 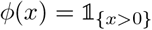 which is 1 when *x* > 0 and zero otherwise. When implemented in the mass action o.d.e., the resulting mass action term contains factors 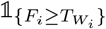 that force reactions to occur only when the concentration of *W_i_* cross thresholds *T_W_i__*. Biologically speaking, this would mean that either of the species *S*_1_ and *S*_2_ is able to take up the excretion products of the other as soon as its concentration is above the threshold.

In the second case, we choose *ϕ* as a continuous sigmoidal activation function with 0 ≤*ϕ*(*x*) ≤1. A continuous activation function with similar behaviour as a discontinuous threshold function is given by *ϕ_ε_*(*x*) = *x*/(2*ε*) + 1/2 for |*x*| ≤*ε*, *ϕ_ε_*(*x*) = 0 for *x* ≤—*ε* and *ϕ_ε_*(*x*) = 1 when *x* ≥ *ε*. A typical such sigmoidal activation function is the Hill function,

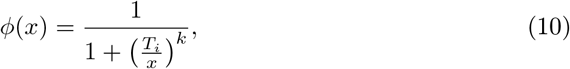

where *k* is the Hill coefficient.

Activation of feedbacks through steep sigmoidal functions are regularly used in description of gene regulatory networks to describe transcriptional activation or repression as a function of transcription factor and effector concentrations (see e.g. [49]). Similarly, uptake of the primary resource R and utilisable waste molecules W can be described by physical contacts and binding of the substances to transporters at the cell surface, justifying the use of a Hill activation function. The final growth model for two species with interaction by cross-feeding for a general activation function *ϕ*, is then described by the following (simplified again by eliminating the intermediate species *P*_1_ and *P*_2_(see [41]). We call this a regulated bacterial network (RBN).

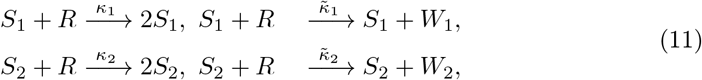

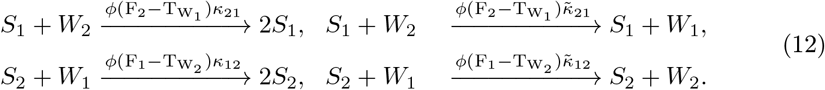

### 3.2 Parametrisation of co-culture growth under substrate competition

#### 3.2.1 Co-culture growth prediction from monoculture-derived kinetic parameters without cross-feeding

We experimentally use a tractable system of two fluorescently labeled bacterial strains *P. veronii* and *P. putida* to test and verify the scenarios of resource competition and resource indifference, under inclusion of cross-feeding or not. Both strains were hereto grown in liquid suspension, either as mono-culture or as a co-culture, with different starting ratios and/or total abundances of the strains. Population growth and relative strain abundances were measured from increases in culture turbidity, strain-specific fluorescence and flow cytometry. We first tested a condition where we expected the strains to be in direct competition, because of utilization of the same substrate (succinate, dissolved at 5 mM). Our first assumption was that single resource competition among two species in co-culture can be described from their inherent individual growth kinetic properties, which can be measured from mono-culture growth.

The relevant growth kinetic parameters extracted from mono-culture growth include the maximum growth rate 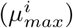, the half velocity constant 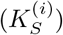 and the biomass yield (*ρ_i_*). These parameters can be expressed as functions of the reaction rates (*κ*_1_, *κ*_2_ and *κ*_3_) (see equation 3, and Appendix S2 to have their explicit form).

First, we estimated the density distributions of the kinetic parameter values from half of the data sets (n = 5) using the Metropolis-Hasting algorithm with a Markov Chain Monte Carlo approach (Fig. 2A and B), and converted these to biomass values using separate empirical cell dry weight estimations (*Materials and methods*). Growth rates of *P. veronii* were 2.2 times slower, and its fitted yields were 75% of those of *P. putida* (Fig. 2A). In contrast, the fitted *K_S_* of *P. veronii* for succinate was lower than that of *P. putida*. The estimated kinetic parameters from half of the replicates led to a proper Monod-simulation of biomass growth of both *P. putida* and *P. veronii*, closely overlapping with the observed increase in biomass in the other five replicate data sets (Fig. 2C). Only exception was a slight decrease observed in optical density values of stationary phase *P. putida*, which the growth model does not take into account. This indicated that we could use the estimated parameter values to simulate co-culture growth.

**Figure 2.**
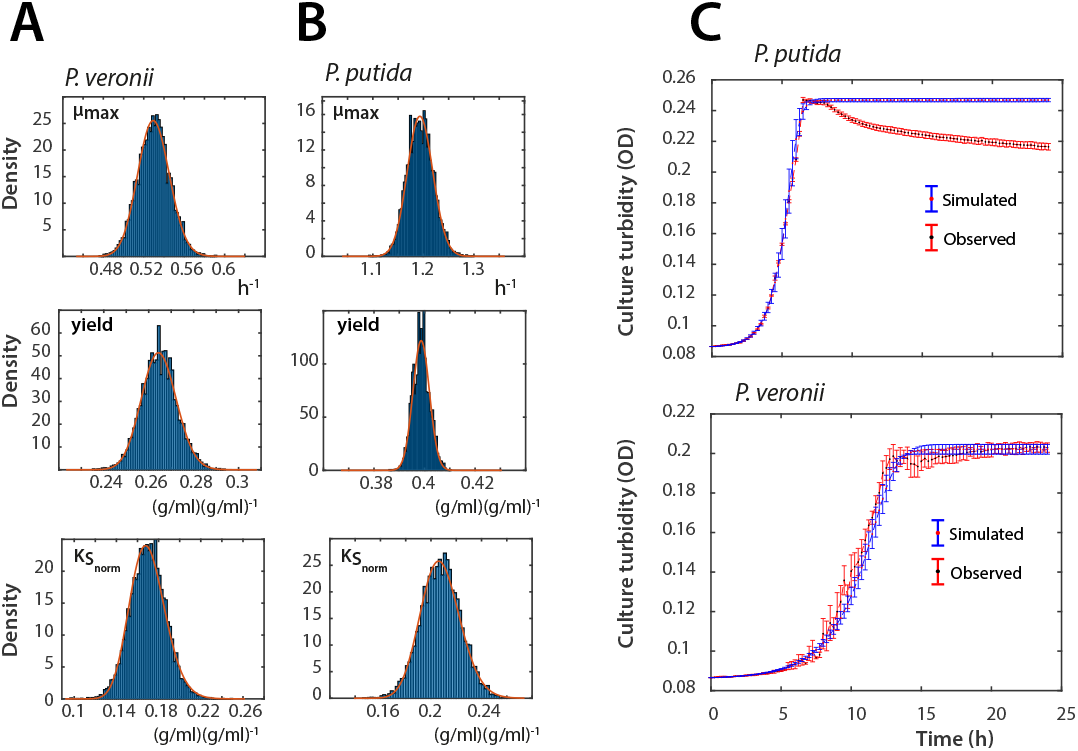
Extracted growth kinetic parameters from mono-cultures of *P. veronii* and *P. putida* using the Monod model. (A) Estimations of *μ_max_,ρ* (yield) and *K_S_* for *P. veronii*. Diagrams show histogram (blue bars) and log-normal (red lines) distributions of parameter estimations from 5 out of 10 biological replicates of *P. veronii* grown in mixed liquid suspension on 5 mM succinate, by using the Metropolis-Hasting algorithm (5.3). Note that 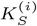 is normalized by dividing by the initial biomass in *g*. (B) as A, but for *P. putida*. (C) Simulated (blue) versus observed growth (red) in mono-cultures of *P. veronii* and *P. putida*, using the parameter sets of A and B. Observed growth is plotted as mean culture turbidity ±one *sd* from the 5 replicate cultures not used for parameter fitting. Culture turbidity transformed from calculated biomass by measured conversion factors in g cell dry weight per ml OD culture (see Materials and methods).

Turbidities of co-cultures with 5 mM succinate increased similarly as those of *P. putida* alone (Fig.3A), suggesting that they consisted in majority of *P. putida*. This was confirmed qualitatively by measurements of the species-specific fluorescent markers during growth, which showed almost similar *P. putida* fluorescence in the co-as in the monoculture, and less than 20% GFP fluorescence from *P. veronii* in the co-culture (Fig.3B). (Since the per cell fluorescence changes as a function of growth phase, the fluorescence plots cannot be converted to species-specific biomass.)

**Figure 3.**
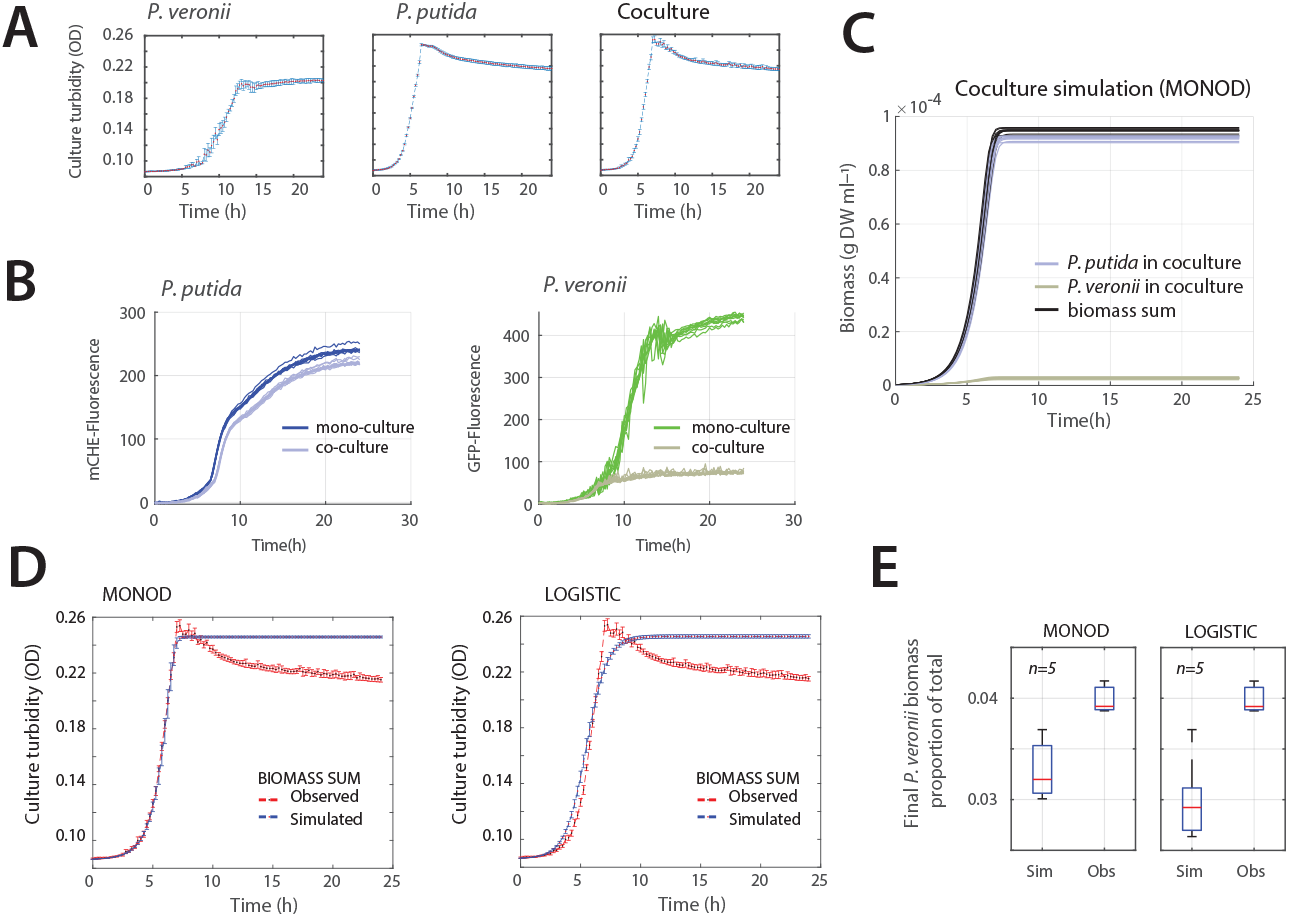
Observed and simulated growth of *P. veronii* and *P. putida* co-cultures under substrate competition. (A) Experimental observations of growth on 5 mM succinate (as culture turbidity). Data points indicate the mean culture density (red dots) ±the calculated standard deviations (vertical error bars, *n*=10 replicates). The co-culture starts as a 1:1 mixture of both species’ cell numbers. (B) Species-specific fluorescence in mono- and co-cultures. *P. putida* read in the mCherry and *P. veronii* in the GFP channel. Lines show individual replicates (*n* = 10). (C) Simulations (*n* = 10) of individual species growth and summed biomass in 1:1 starting co-cultures on 5 mM succinate as sole carbon substrate, using the Monod co-culture model and kinetic parameter sets from mono-culture estimations, without any cross-feeding interactions. (D) Comparison of Monod and logistic co-culture models for simulating the total co-culture biomass growth on 5 mM succinate. (E) Final proportion of *P. veronii* (PVE) of the total biomass for both co-culture models at steady-state (experimental observations use the species-specific cell counts from flow cytometry transformed into species-specific biomass as g DW ml^-1^).

Simulated co-culture growth with kinetic parameter estimates from the individual mono-cultures predicted that the population of *P. veronii* indeed represents a minority in the co-culture (Fig.3C). Prediction of summed co-culture biomass followed closely the observed co-culture growth, but with the Monod co-culture model giving a better fit than the logistic model, which tended to deviate in the early exponential and stationary phase (Fig.3D; for this reason we do not consider the logistic model in further simulations reported here). As expected, both models did not capture the observed decrease in culture turbidity, because neither of them includes any process that would describe biomass decrease. Both models predicted stationary phase proportions of *P. veronii* biomass of approximately 3 % instead of the 4 % measured empirically by flow cytometry in the co-cultures (Fig.3E), n = 10 simulations and n = 3 biological replicates, p-value = 0.0425 using two-sided t-test). This small but statistically significant difference would indicate that the bulk part (75%) of the growth behaviour of both species in co-culture is determined by their inherent kinetic physiological differences, whereas 25% must be due to interactions between *P. veronii* and *P. putida*.

#### 3.2.2 Consistent deviation from predicted competitive behaviour at imbalanced starting cell mixtures

In order to more systematically test the deviation of co-culture simulations with experimental observations, we repeated co-culturing at different starting cell densities and ratios of *P. veronii* to *P. putida*, again using 5 mM succinate as the sole shared carbon and energy source for the cells. Model simulations over a range of starting cell ratios (0.1 —0.999 of *P. veronii* to *P. putida*, Fig. 4A) and/or densities (from 10^5^– 10^7^ cells per ml) suggested that, purely from inherent growth kinetics, *P. putida* will dominate the co-cultures in steady-states, except with 10 - 100 fold surplus of *P. veronii* and high cell starting densities (Fig. 4B). At higher starting cell densities, the availability of resources permits fewer generations of growth, as a result of which *P. putida* cannot gain that much advantage as it would at low starting cell densities (Fig. 4B). By comparing to actual measured species proportions under those conditions, we see that kinetic predictions are in agreement to empirical outcomes only when species ratios are equal at start (1:1, Fig. 4C). In contrast, at imbalanced starting ratios, either of the two species benefits from the presence of the other in improving its growth. For example, at starting ratios of *P. veronii* to *P. putida* of 1:100 and 1:10, the observed final *P. veronii* proportion is statistically significantly higher than expected from its derived mono-culture kinetic parameters, whereas at ratios of 100:1 and 10:1, its final proportion is lower than expected (and *P. putida* proportions are higher; Fig. 4C). This confirmed that the species are likely to engage in some other interaction than only direct competition for resource, as a result of which - in particular, relatively small starting abundances of one species can profit from a larger abundance of the second species.

**Figure 4.**
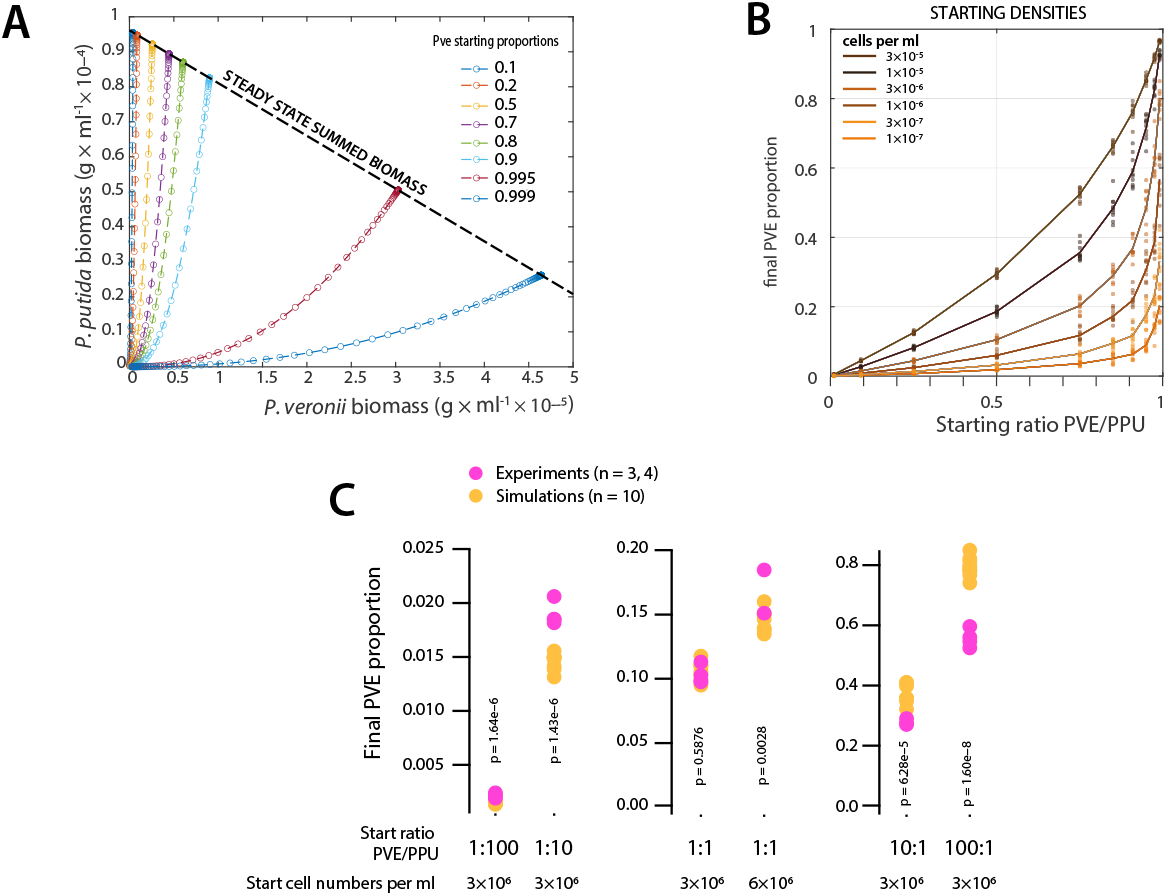
Effect of initial biomass concentration and starting cell ratios in a two-bacterial system growing on a single substrate on their steady-state distributions. (A) Biomass of *P. veronii* and *P. putida* given in g DW per ml with an initial resource (succinate) concentration of 5 mM or 2.4 ×10^-4^g ·ml^-1^. Monod-simulated growth trajectories (without cross-feeding) for the different species at their indicated starting proportions as colored lines and symbols according to the legend. (B) Steady-state biomass concentrations as a function of different starting cell ratios and starting cell densities (colors, in g biomass DW per ml). Symbols correspond to individual replicate simulations, with log-random sampled kinetic parameters from estimated distributions of Fig. 2. (C) Difference of observed (magenta) versus simulated (ochre) final *P. veronii* proportions at different initial ratios (bottom) and two starting densities. p-values stem from paired two-sided t-test comparisons.

#### 3.2.3 Cross-feeding models suggest utilization of part of excreted metabolites at imbalanced starting cell ratios

To estimate the potential effect resulting from cross-feeding interactions as we theorised conceptually above in the species reaction models, we simulated the production of waste (the W-term in equation 3 and 4) from the fitted *κ*_1_, *κ*_2_ and *κ*_3_ values for both of the species in mono-culture. In absence of allowed cross-feeding (Fig. 5A), one can see that both species convert a significant quantity (60%) of the primary substrate succinate (5 mM, equivalent to 2.5 ×10^-4^g C per ml) into waste (here taken as the sum of all product not incorporated into biomass, including CO_2_). This is not surprising, as excretion of up to 50% of re-usable carbon metabolites (apart from released CO_2_) has been routinely described to occur from individual strains growing in batch culture with large primary substrate quantities (i.e., mM-range) [40,47]. By allowing cross-feeding to occur, one can observe that, depending on the cross-feeding model, the appearance of waste in the co-culture is diminished and profiting to biomass growth of the reciprocal species (as expected; Fig. 5B-D). This simulation already visually indicates that allowing all waste to be re-utilized by the reciprocal species (as in model B of equation 7) is not in agreement with observed co-culture growth, because it leads to extended growth at later phases (i.e., > 10 h; Fig. 5B, compare to Fig. 3). Interestingly, the models further predict that it is primarily *P. putida* which at low relative starting abundances can profit from waste excreted by the larger thriving population of *P. veronii*, but less so in reverse (Fig. 5C and D). By comparing the steady-state values of the simulated *P. veronii* waste concentration at the highest starting ratio of Pve:Ppu= 100: 1 in absence of cross-feeding and with threshold cross-feeding, we would obtain a total of 1 ×10^-4^g C per ml that is excreted by *P. veronii* and used by *P. putida* for its growth (Fig. 5C and D). Finally, this simulation also shows that cross-feeding is most optimally observed at unequal starting cell ratios.

**Figure 5.**
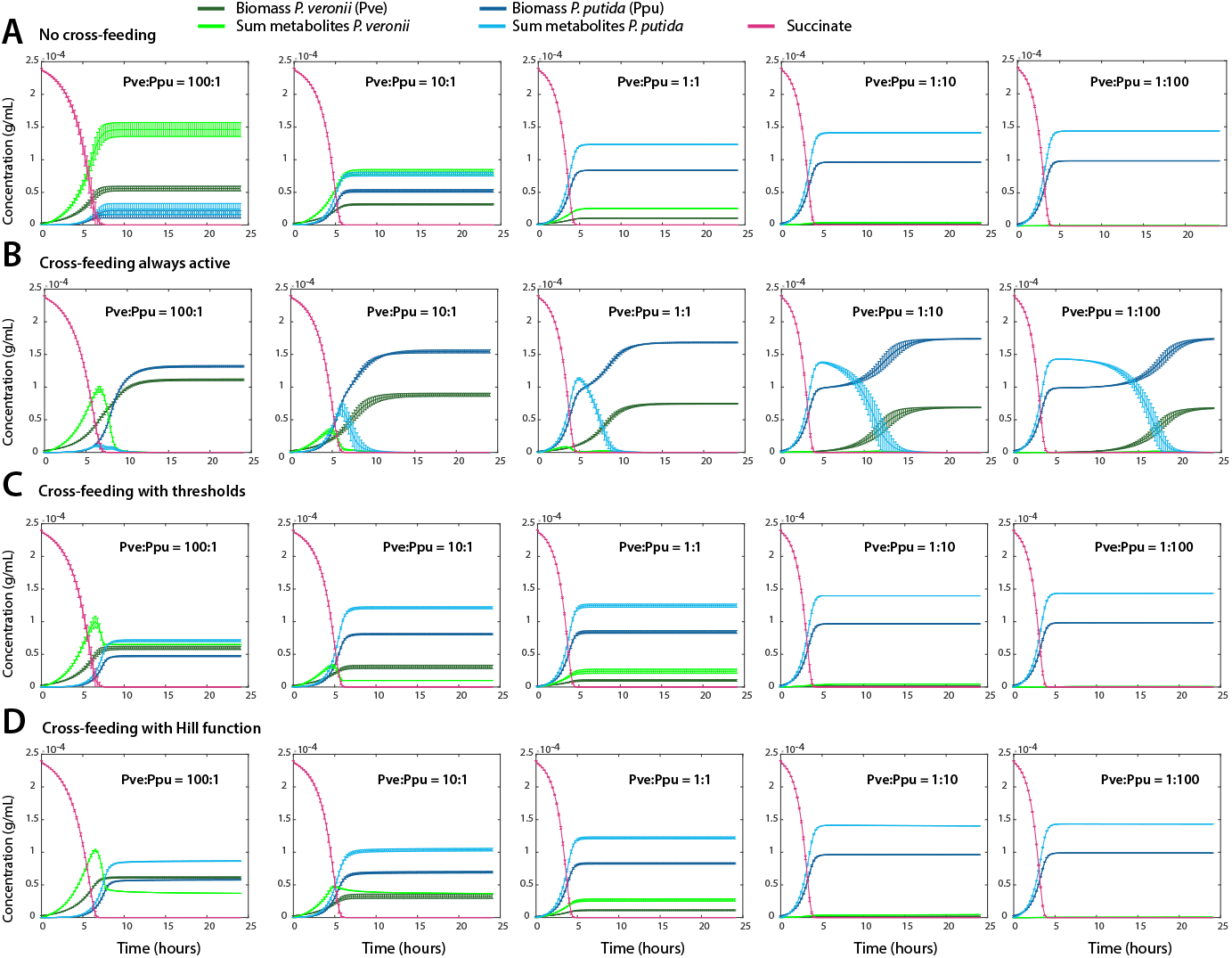
Simulated biomass growth and waste formation in co-culture of the two species, with or without waste re-utilisation. A-D. Biomass growth of either species *P. putida* (red) or *P. veronii* (blue) on a single shared carbon substrate (5 mM succinate), and predicted waste concentrations (light and dark green), for five different cell starting ratios (100:1, 10:1, 1:1, 1:10 and 1:100, as indicated), and 1 ×10^6^ cells per ml at start. Simulations in (A) assume no cross-feeding (Monod model equation 4), (B) permanent cross-feeding of appearing waste, according to equation 7,8. Simulations in (C) follow cross-feeding with the discontinuous threshold function (equation 11) with threshold values (in [g/mL]), from left to right, *T*_*W*_1__ = 6.5 ×10^-5^, 1.0 ×10^-5^, 2.8 ×10^-5^, 2.8 ×10^-5^, and 2.8 ×10^-5^; *T*_*W*2_ = 1.4 ×10^-4^, 1.4 ×10^-4^, 1.4 ×10^-4^, 1.4 ×10^-4^, and 1.43 ×10^-4^. Model in (D) assumes cross-feeding following a Hill activation function (equation 12) with parameter values, *T*_*W*_1__ = 4.73 ×10^-5^[g/mL], *T*_W_2__ = 1.73 ×10^-4^[g/mL] and *k* = 34. *T*_*W*_1__ corresponds to waste produced by *P. veronii* that is used by *P. putida*; and *T*_*W*_2__ vice versa. Plots show the means (lines) from *n* = 10 simulations with error bars representing ±one standard deviation. Variation is introduced from resampling the distribution of kinetic parameters as in Figure 2.

In order to better understand the effects of thresholding parameters, we simulated a range of threshold values in the discontinuous function of equation 9 on the predicted stationary phase abundances of *P.veronii* and *P.putida* in co-culture, at a starting ratio of Pve:Ppu of 10: 1. In case of the discontinuous threshold function, the uptake rates of byproducts are assumed to be equal to that of the primary resource (hence, rates *κ*_1_4__ and *κ*_2_4__ equal to *κ*_1_1__ and *κ*_2_1__, respectively, as in eq.4). In this case, only the value of the threshold parameters *T*_*W*_2__ and *T*_*W*_1__ impacts the predictions of the stationary phase species biomass proportions. As one can see in Fig. 6A-C), varying the threshold values on the utilisation of *W*_1_ by *P. putida* and of *W*_2_ by *P. veronii* has an important effect on their stationary phase proportions; their closeness to experimental observations and the corresponding utilization of reciprocal waste products. For instance, the steady-state proportions of *P. veronii* are lowest when its threshold for using the byproducts from *P. putida* is highest (i.e., at *T*_*W*_2__= 0.00015, Fig. 6A), and *P. putida* continuously takes away byproducts from *P. veronii* (i.e., *T*_*W*_1__= 0). On the contrary, steady state proportions of *P. veronii* are highest when *P. putida* has a high threshold on using byproducts from *P. veronii* and *P. veronii* has a low threshold on using byproducts from *P. putida* (i.e., *T*_*W*_1__ is 0.000135, and *T*_*W*_2__ is 0, Fig. 6A). The latter scenario is less likely to be in agreement with experimental data, as plotting the corresponding ratio of simulated versus observed stationary phase species ratios shows (Fig. 6B). This suggests that *P. putida* is more efficient in utilizing byproducts from *P. veronii* than the other way around, and, as mentioned, is predicted to profit from up to 1 ×10^-4^g C per ml waste product from *P. veronii* (Fig. 6C). The physiological reason for this efficiency could lay in the lower *K_m_* of its uptake systems or in the types of metabolites produced and released by *P. veronii*.

**Figure 6.**
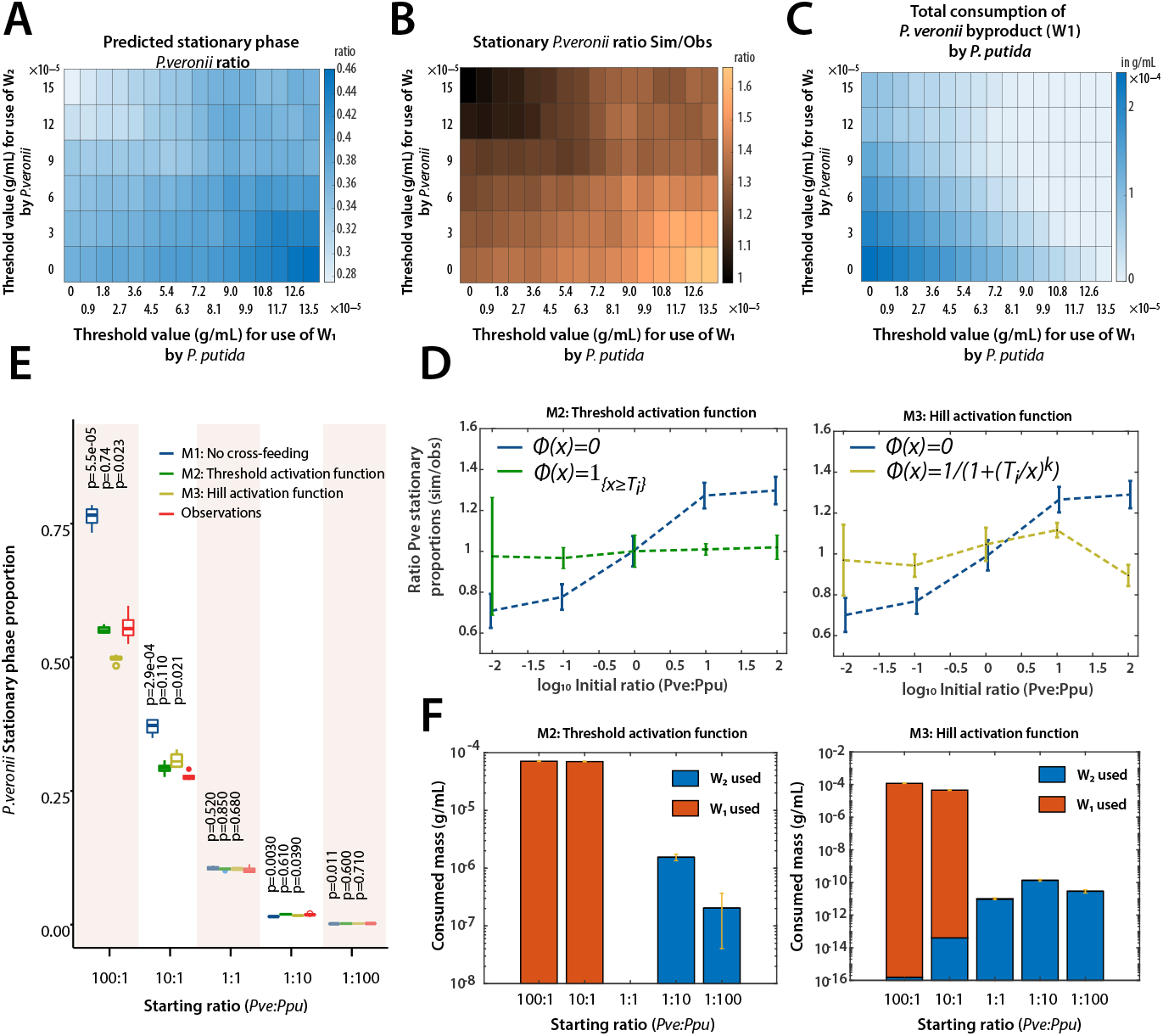
Effect of thresholding on cross-feeding between both species in co-culture. (A) Predicted stationary phase ratios of *P. veronii* biomass as a function of the discontinuous threshold value for activation of cross-feeding by *P. putida* on the byproducts of *P. veronii* (*W*_1_), and vice-versa (*W*_2_). Ratios plotted as heat map according to the color legend as indicated. (B) Similarity of simulated versus observed stationary phase biomass proportion of *P. veronii*, taken as the ratio; again as function of selected values of *T*_*W*_1__ and *T*_*W*_2__. (C) Predicted waste utilization by *P. putida* as a function of of *T*_*W*_1__ and *T*_*W*_2__. (D) Mean relative difference of simulated versus observed stationary phase proportions of *P. veronii* in co-culture with *P. putida* on 5 mM succinate at different starting cell ratios (as indicated 100: 1, 10: 1, 1:1, 1: 10, and 1: 100). Three model predictions are shown: M1, without cross-feeding (in blue), overlaid with either M2 (Model with discontinuous threshold function, in green); or with M3 (Hill activation function, in yellow). (E) Comparison of observed steady-state *P. veronii* biomass proportions (observations, in red; *n =* 4 replicates) at different starting cell ratios, to those resulting from the three model simulations (M1, M2 and M3; *n =* 4 replicates), as above. Box plots show the median, second and third quadrants, and lines indicating the 25th and 75th percentiles. P-values calculated by t-test in comparison to the empirical values. (F) Total estimated utilization of waste (in g *C/mL*) for each of the co-cultures and varying initial cell ratios, as indicated. In blue, consumption of byproducts by *P.veronii*. In red, consumption of byproducts by *P.putida*. Model M1 corresponds to equation 4; model M2 to the discontinuous threshold model of equation 9, and M3 to the Hill activation function for waste utilisation of equation 10. Steady-state cell concentrations of both species were measured by flow cytometry and identified on the basis of the respective fluorescence marker; and converted to biomass using calculated single cell mass values, as described in the Materials section. Model M2 simulations used threshold values of *T*_*W*_2__=1.4 ×10^-4^, and *T*_*W*_1__=6.5 ×10^-5^(ratio 100:1); 1.0 ×10^-5^(ratio 10:1); and 2.8 ×10^-5^(all other ratios). Model M3 simulations used the fitted parameters: *T*_*W*_2__= 1.73 ×10^-4^, *T*_*W*_1__ = 4.73 ×10^-5^, and *k* = 34.5.

Comparison across different simulations (variations from subsampled kinetic parameters), the five starting ratios and the three models, showed that thresholding gives the better explanation of the observed steady-state *P. veronii* proportions in co-culture with *P. putida* on 5 mM succinate (Fig. 6D and E). Threshold values for the discontinuous model (M2) were derived from the parameter screens, but are arbitrary fits for each of the starting cell ratios. In contrast, the Hill function parameter values (eq: 10) were fitted on the experimental data for the complete data sets (as described in the Materials section). Using the fitted values, we find a maximal difference of about 13% (Fig. 6D; M3) at stationary phase between the measured and simulated abundances. In comparison, the model without cross-feeding predicts a maximum difference of about 30%. This indicates that there is a significant interspecific interaction term that develops during growth on a single shared resource, and from which the lesser abundant species can profit. This is counterintuitive from the concept of competition but makes sense from microbiological perspective, because an excreted waste by an abundant population contains potentially sufficient carbon to support measurable growth of a small population of a second species, but not of a large one. However, both thresholding models on cross-feeding predict that *P. putida* can utilize 2 (model M2) to 6 (model M3) orders of magnitude more of the reciprocal waste than *P. veronii* (Fig. 6F).

### 3.3 Substrate indifference: Co-culture growth with two independent resources

To contrast direct substrate competition with appearing cross-feeding with a situation in which both species would be solely driven by growth kinetic differences, we designed an experimental scenario of substrate ‘indifference’. We repeated co-culturing of *P. putida* and *P. veronii*, but in this case with dual substrates, one of which (D-mannitol) being specific for *P. veronii*, and the other (putrescine) for *P. putida*. Indeed, we could not measure growth of *P. putida* on D-mannitol, whereas *P. veronii* developed to one-fifth of its biomass on putrescine in comparison to D-mannitol, albeit with relatively slow growth (Fig. 7A). However, in a 1:1 starting ratio co-culture on D-mannitol or putrescine, there was no distinguishable growth of *P. putida* or *P. veronii*, respectively, indicating effective unique and exclusive primary substrates to each of the species (Fig. 7A). In contrast, the culture turbidity of the co-culture on putrescine was lower than that of *P. putida* alone on putrescine (Fig. 7A). Co-cultures with both substrates simultaneously and with varying species’ starting ratios (from 100: 1 to 1: 100, as before) showed growth of both species (judged from their specific fluorescence marker) to approximately the same final levels, independent of their starting ratio (Fig. 7B). The apparent (visible) difference in onset of growth of either species in the co-culture is due to its varying population size in the starting mixtures, which is more clearly explained in the simulated Monod growth using fitted kinetic parameters in absence of any assumed cross-feeding, as in equation (5) (Fig. 7C; the species-respective *κ*_1_, *κ*_2_ and *κ*_3_ again deduced from the measured *μ_max_, K_S_* and yields as shown in Fig. 2).

Since there was no competition for the resource, we hypothesised that the population growth of both species would be independent of any appearing byproducts, in contrast to the case with succinate as unique (competitive) resource, and should reach the same abundances in co-culture as in monoculture. Indeed, both species grew in parallel in the co-cultures with both substrates (Fig. 7C), with a mean stationary proportion of *P.veronii* across all starting ratios of 54.0% (±one SD of 3%) (Fig. 7D). This is in contrast to the case of succinate, where the proportions of *P. veronii* varied between 0.2 and 60% (Fig. 6E). In addition, one can observe how the community growth is almost a ‘sum’ of two overlaying independent species growth curves, which become more delayed, the smaller is the starting proportion of *P. veronii* (Fig. 7C). The difference in the extension of the co-culture growth phase is due to the faster maximum growth rate of *P. putida* on putrescine (1.54 h^-1^) than *P. veronii* on D-mannitol (0.36 h^-1^) (Supplementary Fig. 1).

Although the observations in this case were well explained by the growth kinetics alone of either species (i.e., without inclusion of cross-feeding), we still observed small deviations between the simulated (predicted) and observed stationary phase abundances of *P. veronii* (Fig. 7D). However, in this case it was the observed *P. veronii* stationary phase proportion that increased from 51.1% for a starting ratio of (Pve:Ppu)= (100: 1), to 57.6% for a starting ratio of = (1: 100), whereas the simulations predicted a constant ratio of 54%. Interestingly, the measured stationary phase abundance of *P. putida* cells remained the same among all starting ratios (2.4 ±0.06 ×10^9^cells per ml, equivalent to a biomass of 1.69 ×10^-4^±4.6 ×10^-6^[g/mL]). If at the same time the final measured cell number of *P. veronii* increased from 9.8 ×10^8^at a Pve:Ppu starting ratio of 100: 1 to 1.4 ×10^9^ cells per ml at ratio of 1: 100, this could mean that either the *P. veronii* cell size decreases or that more of the available resources is converted into biomass.

Despite both exclusive substrates, simulations thus again favor some cross-feeding, but in this case by *P.veronii* of byproducts coming from *P. putida*, following equation (5). Cross-feeding can again be described by a discontinuous threshold under constant *P. veronii* yield of 34.42% (Fig.7 D). In contrast, the biomass of *P.putida* is not impacted by the cross-feeding, and its yield of 34% remains constant in models with and without cross-feeding. Allowing cross-feeding to occur would then lead to utilization of byproducts as visualized in Fig.7 E and Supplementary figure 2.

## 4 Discussion

We developed a generalized cross-feeding model for two-species interactions, which links appearing metabolites in the extracellular medium to changes in growth kinetic parameters. Cross-feeding was best explained as an activation threshold function, which could be experimentally parameterized under two interaction scenarios: (i) primary single substrate competition and (ii) substrate indifference. Comparisons of model simulations to experimental data showed that modeling with inherent Monod kinetic parameters without interspecific interaction terms are insufficient to explain co-culture growth. This was particularly true for cases where the initial abundances of both species were unequal, and the numerical minority could proliferate more than expected. From an ecological perspective this is an important notion, because it would allow minority species to grow and survive better than expected from pair-wise competition assays that are conducted at equal species-ratios. We further showed that the model holds for both the situation of imposed primary substrate competition and the substrate indifference, albeit it with reversal of the species profiting most from the metabolite cross-feeding.

Our mathematical description extended generalized consumer-resource models [17] with interspecific interactions from chemical reaction network CRN [20], which is similar to propositions as in Ref. [40] that assume cross-feeding mechanisms from metabolic generalists to other species on their metabolic by-products. Although these authors experimentally proved that such cross-feeding mechanisms are necessary to ensure observed species coexistence in microbial communities through collective interactions, and that metabolite concentrations are high enough to ensure growth of other species, their proposed permanent cross-feeding loops [40] cannot mathematically hold. As we demonstrate in Proposition 1, permanent cross-feeding loops would lead to vanishing equilibrium waste concentrations and biphasic co-culture growth, which we did not observe empirically.

As an alternative, we introduced a regulatory mechanism that allows for activation of the cross-feeding positive feedback loop only above a threshold concentration, which overall gave better predictions of experimentally observed co-culture growth in a mean-field resource-limited environment (such as batch liquid suspended cell culture).

Regulated interaction networks are commonly to describe for gene regulation, where (positive and negative) feedback loops can be active (ON) or inactive (OFF) depending on the concentration of transcription factors or inducers and effectors. For example, during transcription, which is inherently noisy, promoters switch randomly between ON and OFF states. The switching rate *ϕ(F*) depends in a nonlinear way on transcription factor concentration F, which is properly described by a Hill activation function *ϕ*. The reader can consult, e.g., [50] or [51], [49] for mathematical results on stochastic and deterministic systems involving such ON-OFF switching. We propose here that, similarly, activation functions can be used to describe interspecific growth interactions resulting from excreted/shared metabolites during primary growth.

**Figure 7.**
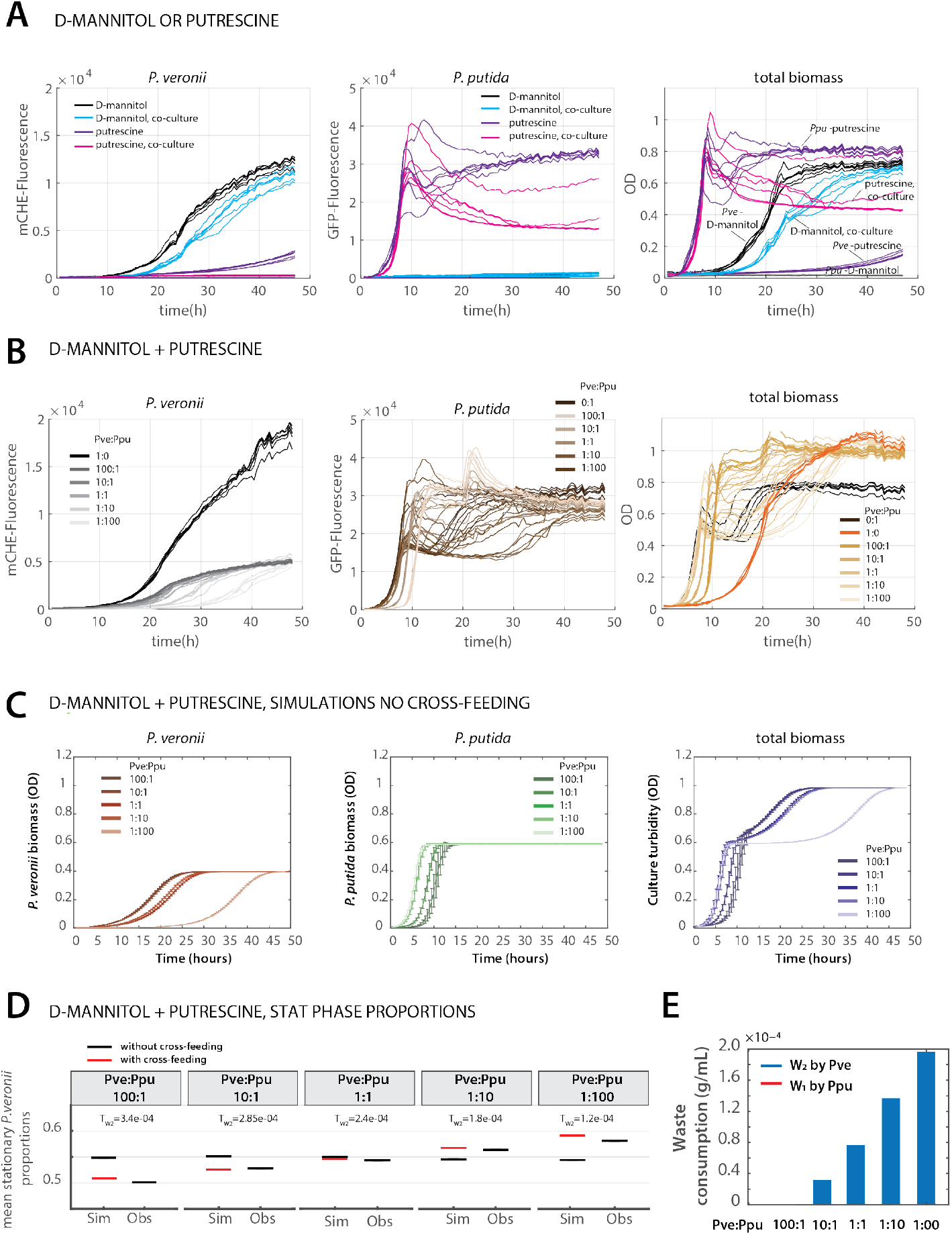
Dual exclusive substrates in co-culture lead to independent growth of each species and inversion of cross-feeding. (A) Growth of *P. putida* or *P. veronii* alone and in 1:1 co-culture with either putrescine or D-mannitol. Graphs show development of mCherry fluorescence (specific for *P. veronii*), of GFP fluorescence (specific for *P. veronii*) and of culture turbidity (n= 6 replicates). (B) Growth of *P. putida* and *P. veronii* alone or in co-culture with mixture of 10 mM putrescine and 6.7 mM D-mannitol, at varying starting cell ratios (as indicated, n= 6 replicates). (C) Simulated growth of *P.veronii* and *P.putida*) under the same conditions and starting ratios as in B, using fitted kinetic parameters in absence of assumed cross-feeding. Data points show means from 8 replicates (points) ±their calculated standard deviation (bars). (D) Mean simulated (Sim) and measured (Obs, from flow cytometry) stationary phase proportions of *P.veronii* in the co-cultures, in absence (black) or presence (red) of assumed cross-feeding (discontinuous imposed threshold values indicated). Variation of the data is smaller than the line size. (E) Calculated utilization of waste byproducts by *P. veronii* and *P. putida* for the threshold-imposed cross-feeding scenarios in panel D.

The difficulty in modeling community growth under inclusion of cross-feeding interactions is that, first of all, maximum growth rates are not known for many of the constituting taxa; neither are their substrate dependencies, nor their capacities and rates to utilise metabolites appearing during growth of other neighbouring taxa. To make more inclusive models, one could expand the simple ‘summed’ waste concentration as we proposed here, by matrices that would cover all (major) individual relevant substrates and products, with measured or estimated rate constants; for example, as proposed in [40]. One could imagine that characterizing the single species substrate and metabolite ‘landscape’ will be facilitated in the future from detailed genome-scale metabolic models, which can also put boundaries on possible growth and reaction rates [13, 52, 53]. Alternatively, one could use lumped rate constants inferred from mono- and co-culture growth data, such as utilised here and proposed elsewhere [14]. Although we did not specifically measure metabolite concentrations appearing during mono- and co-culture growth, there is ample evidence for re-usable byproducts to appear and in the concentration ranges [44–48] that we infer here from kinetic parameter fitting (see, e.g., 4). In addition, we estimate that approximately 25% of the yield in imbalanced starting ratios may originate from cross-feeding mechanisms.

We rigorously confronted and validated the model’s predictions with experimental data for a two-species ecological network. Our experimental results were obtained in a suspended growth liquid environment, which leads to a mean field mathematical model. It will be interesting now to extend to controlled experiments with microbial communities and metabolic networks comprising more than two species, in order to assess the general usefulness of the regulated bacterial network models for predictions of community growth and its corresponding taxa composition. Furthermore, it is relatively straightforward to expand the mathematical framework to spatial situations with multiple species, growing at the expense of diffusible substrates, which will get us closer to understanding and predicting spatially structured communities, see e.g. [17].

## 5 Materials and methods

### 5.1 Bacteria cultivation

*Pseudomonas veronii* 1YdBTEX2 (PVE, strain 5336) is a mini-Tn5 inserted tagged variant constitutively expressing GFP from the P_*circ*_ promoter of ICEclc [54], [55]. *Pseudomonas putida* F1 strain 5789 (Ppu) is a mini-Tn5 tagged variant of the wild-type strain [56] carrying a single-copy *mcherry* gene under control of P_*tac*_. Tagging of F1 was accomplished by conjugation of the corresponding mini-Tn5 construct from *E. coli* DH5a-lpir with *E. coli* HB101 as helper for conjugation. Glycerol stocks of Pve and Ppu from —80°C were plated on nutrient agar containing gentamycin at 10 mg L^-1^ for selection of the mini-Tn5 constructs. Single freshly grown colonies were transferred to liquid 21C minimal media (MM) [57] in glass flasks with 5 mM succinate, and incubated at 30°C at 180 rpm for 16-20 h. To prepare cells for competition experiments, culture samples (10 ml) were centrifuged at 5000 rpm in a F-34-6-38 rotor in a 5804R centrifuge (Eppendorf AG) for 10 min at room temperature. The supernatant was discarded and the cell pellet was resuspended in 10 ml of minimal media salts (MMS, containing, per litre: 1 g NH4Cl, 3.49 g Na2HPO4·2H2O, 2.77 g KH2PO4, pH 6.8). The cell density in these suspensions was measured by flow cytometry (see below), after which the suspensions were used for the competition experiments.

Washed cultures were freshly diluted in liquid medium with 5 mM of succinate, or with 10 mM D-mannitol (for *P. veronii*), or 6.7 mM putrescine (for *P. putida*) as sole added carbon and energy source, either as individual monoculture or as binary mixture, each in ten replicates. Individual (mono-)cultures started with 10^6^ cells ml^-1^. In the co-cultures, the initial ratio between *P. veronii* and *P. putida* was varied from 100: 1,10: 1,1:1,1: 10 and 1: 100, each time starting with a mixture that contained 10^6^ cells ml^-1^. In case of the substrate indifference experiment, we relied on diluting the starting cell suspensions to a culture turbidity at 600 nm (OD600) of 0.05 for *P. veronii* and 0.02 for *P. putida*. Cultures were incubated at 28-30 °C and growth was followed during 24-48 hours by automated reading of the culture turbidity and of GFP/mCherry fluorescence, every 15 (for succinate) or 30 (for D-mannitol and putrescine) minutes in a 96—well plate reader. After 24 (for succinate) and 48 h (for D-mannitol and putrescine), three replicates of each mono-culture and of the co-culture were sampled, diluted and measured by flow cytometry to obtain the exact cell density. These measurements were used to determine a global conversion factor between the number of cells and the culture turbidity (‘optical density’, OD), while acknowledging that this doesn’t take cell size changes into account. Measured culture densities were corrected for the optical density of the medium itself.

#### 5.1.1 Flow cytometry

The cell density in the washed cultures was quantified either using a BD LSRFortessaTM (BD Biosciences, Allschwil, Switzerland) or a Novocyte flow cytometer (ACEA Biosciences, USA). For quantification using the BD LSRFortessaTM flow cytometer, an aliquot of 50 *μ*L of bacterial culture was mixed with 2 mL sterile saline (9 g NaCl L^-1^) solution in a 5 mL polystyrene tube (Falcon, Corning, NY, USA). The following parameter settings were used for cell quantification: FSC = default, SSC = default, FITC = 650 V, PE-Texas-Red = 650 V, injection flow rate = 35 *μ*L min^-1^, and acquisition time = 60 s. Four different FSC-H thresholds were tested for quantification: 400, 600, 800, and 1000. Pve was quantified on its FITC-signal (GFP), Ppu on the PE-Texas-Red signal (mCherry). For detection and quantification using the Novocyte flow cytometer (ACEA Biosciences, USA), washed liquid cultures were diluted 100-1000 times in MMS and 20 *μ*L were aspired at 66 *μ*l min^-1^. Ppu cells were identified at an FSC-H threshold above 500, SSC-H above 150, and a PE-Texas Red-H signal gated to above an empty control (typically: 1000; channel voltage set at 592 V, representative for the mCherry fluorescence). Pve cells were identified at same FSC-H and SSC-H thresholds but on the basis of an FITC-H signal above the empty control (typically, 600; representative for GFP fluorescence; channel voltage at 441 V).

#### 5.1.2 Liquid culture competition experiments

To test competition between Pve and Ppu in liquid culture, we inoculated strains either individually or in combination into MM medium with either succinate (competitive scenario) or a combination of D-mannitol and putrescine as carbon substrates (indifference scenario). For succinate we used 5 mL MM medium with 10 mM succinate in a 29-mL sterile capped glass vial, with access to ambient air, and incubated cultures at 30°C at 180 rpm. The cultures were sampled after 1, 24, 48, 72, and 144 h by removing a 50 *μ*l aliquot, which was diluted with 2 ml of sterile water, to quantify the cell density of either Pve or Ppu using flow cytometry.

Yields from cell numbers and biomass were compared through measured cell volumes with succinate, D-mannitol or putrescine as substrates. The cell dimensions were quantified from phase-contrast microscopy images of culture samples using Fiji [58]. On succinate, the average cell volume of Pve amounted to 1.8 *μ*m^3^ and that of Ppu to 0.8 *μ*m^3^. This was converted to a per cell carbon mass using the conversion factor of 0.264 pg C for an average *E. coli* cell volume of 2 *μ*m^3^[59]. Cell and biomass yields were further inferred from weighed 70°C-dried filtered samples from flow-cytometry counted stationary phase cultures grown on 10 mM succinate. For *P. veronii* this corresponded to 0.38 mg and 8.9 ·10^8^ cells per OD; for *P. putida* to 0.46 mg and 9.5 ·10^8^ cells per OD.

### 5.2 CRN and mass action kinetics

#### 5.2.1 Modeling with CRN

Our starting modeling framework was the logistic model, which assumes that growth can be described as a reaction involving a species *S* that enters in contact with a resource *R*, which it transforms to new cell and duplicates. This process has been described by the reactions (1)

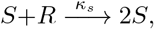

which describes the consumption of an item of *R* by an item of consumer *S*, which results in the creation of an additional item from *S*, and (2)

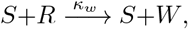

which describes the reaction where the consumption of an item from *R* by an item from *S* produces an item from *W*. We then introduced the kinetic rates 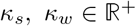 and the concentration or abundance per unit of volume *X* = [*S*], *C* = [*R*], *F* = [*W*] of the species *S*, the resource *R* and the waste *W*, respectively. We claim that the related mass action o.d.e. is given by

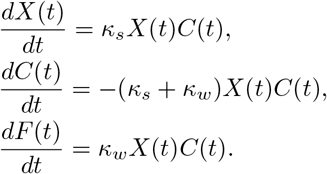

One can move from chemical reactions like (1,2) which describe random consumption and creation of molecules (integers), to a continuous deterministic mass action o.d.e because of the law of large numbers: in this special case, the total number *N* of molecules is preserved, and assuming a well-mixed environment, one can show that the concentration abundances converge in probability as *N* →∞toward the solution of the mass action o.d.e., see, e.g. [60].

#### 5.2.2 Mass action kinetics

Modelling with CRN allows the use of both stochastic and deterministic tools, see, e.g., [49]. We follow [61] which provides a clear exposition of the basic tools and results from deterministic CRN theory. Consider *s* species *S*_1_,…,*S_s_*, *s* ≥ 1, whose interactions are described by r reactions, denoted symbolically by reactions schemes

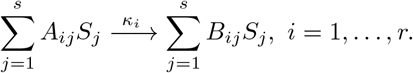

where the coefficients *A_ij_* and *B_ij_* are stoichiometric coefficients which are non-negative integers. One can use *r* x *s* matrices *A* =(*A_ij_*) and *B* =(*B_ij_*) to write the above reactions schemes in a more compact way as

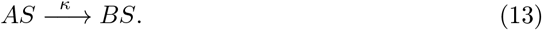

where *S* =(*S*_1_,…, *S_s_*)^*T*^ is the column vector of species and *κ*= (*κ*_1_,…, *κ_r_*)^*T*^ is the positive vector of kinetic rates. Given a vector *X* = (*X*_1_,…, *X_s_*)^*T*^ and a non-negative *r* x *s* matrix *A*, let *X^A^* be the *r* dimensional vector whose ith entry is the product 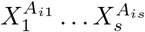. The mass action o.d.e. associated to the reaction (13) is

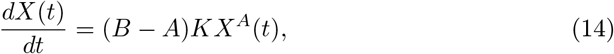

where *K* is the diagonal matrix having the kinetic rates on the diagonal.

Importantly, one can prove (see, e.g., [61]), that, starting from non-negative initial conditions *X_j_* (0) ≥ 0, the solutions to the o.d.e. (14) remain non-negative for all times.

#### 5.2.3 Generalized consumer-resource models

We present here the generalized consumer-resource model described in [40] and [62]. Let *W_k_, k* = 1,…, *M* be the byproducts or waste species and let *S_i_, i* = 1,…, *n*, be the set of consumers. Let *c_ik_* and *l_k_* be the rate at which species *S_i_* uptakes resource *W_k_* and the leakage factor for resource *k* with 0 ≤ *l_k_* ≤1. Let *σ* be a single-valued function encoding the relationship between resource supply levels and uptake rates. Typically, *σ* is linear or a sigmoidal function of Hill type. Let *m_i_* be the minimal energy required for growth of species *S_i_*. The biomass of species *S_i_* at time *t* is denoted by *N_i_* (*t*), and the abundance of waste *W_k_* is denoted by *R_k_*(*t*). The o.d.e. describing the time evolution of *N_i_* is

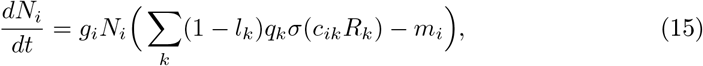

where the function *q* =(*q_k_*) might be, for each resource *W_k_*, the maximal ATP yield of resource *W_k_*, and *g_i_* is a positive proportionality factor. Concerning the time evolution of resources, let, for each species *S_i_*, the stoichiometric matrix *D* = (*D_kl_*)_1≤*k,l*≤*M*_ encode the number of molecules of resource *k* secreted to the environment per molecule of resource *l* taken up. The secretion matrix *P* = *D^T^* is assumed to be stochastic.

Within this modelling framework, the production rate of resource *W_k_* from secretions is

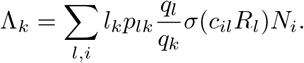

The o.d.e. governing the time evolution of resources is then

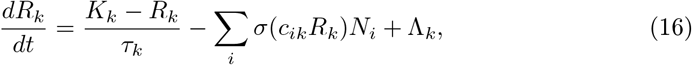

where *K_k_* is the initial abundance of resource *W_k_*, and *τ_k_* is the transfer rate during batch culture passaging. We assume here that *τ_k_* >> 1, so that the first term (*K_k_*—*R_k_*)/*τ_k_* can be neglected.

**Example:**

Consider the CRN (7,8)

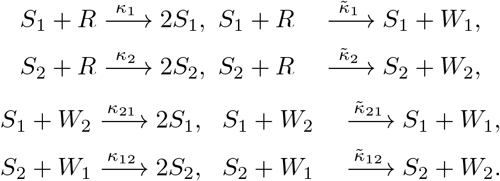

of mass action kinetics o.d.e.

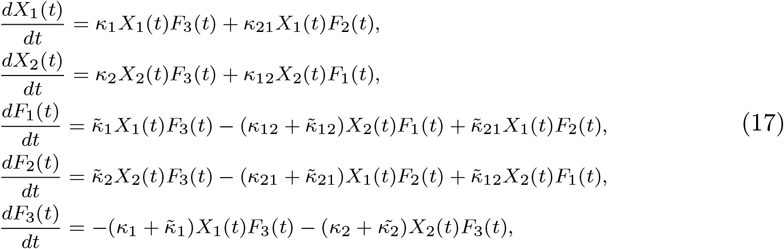

where *X_i_* are the concentrations of species *S_i_, i* = 1,2, and where *F_k_*, *k* = 1, 2, 3 are the concentrations of waste *W_k_, k* = 1, 2 and of resource *R*.

This o.d.e. is similar to that defined in (15,16). Appendix S4 considers these two systems of differential equations and arrives at the non-identifiability of generalized consumer-resource models within our framework. One can find a one dimensional manifold of parameters *g_i_*, and *l_k_, q_k_, p_lk_* and *c_ik_* so that (15,16) and (17) coincide for given parameters 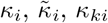 and 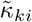.

### 5.3 Models and simulations

We assume that the d-dimensional o.d.e. which models species population growth is of the form

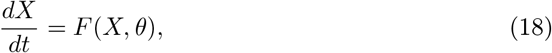

where the unknown parameter 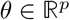 contains the reaction rate constants of the underlying chemical reaction network. So, given a parameter **Θ**, one can simulate the solution of the o.d.e. for some given initial condition to get *X*(*t*, θ), *t* ≥ 0. Then, the value of the resulting solution is compared to the observed experimental values *y_n_*, for time points *t_n_, n* = 1,…, *N*. The statistical model includes noise terms and is of the form

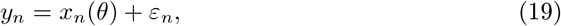

where *x_n_*(*θ*) = *X* (*t_n_, θ*) and the *ε_n_* model experimental errors. This process is replicated *R* times, each simulation and experimental replicates involving *N_r_* time points. The resulting replicates are denoted by *y_r,n_,x_r,n_*(*θ*) and *ε_r,n_, r* = 1,…, *R*. We set for convenience *y_r_* = (*y_r,n_*)_*r*=1,…,*N_r_*_.

Reaction rates constants for *P. veronii* and *P. putida* were extracted from mono-culture growth using the Metropolis-Hasting algorithm with Markov Chain Monte Carlo approach to estimate their probability distributions. The advantage of this procedure is that it estimates the variation in the growth rates, which can be included in simulations of the experimental results. For this, we separated the complete dataset (*n* = 10 replicates) in two halves. On one, we estimated growth rates and the remaining data were used to compare simulations and empirical observations.

The Metropolis-Hasting algorithm seeks to estimate the conditional probability distribution of a model parameter 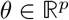 that generated a set of empirical observations 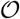. Using Bayes’ theorem and the information we have from the observations generated by *θ*, we can obtain the posterior probability function of the model parameter,

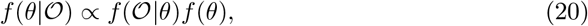

where 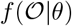 is the information (likelihood) of the data generated by *θ*, and *f* (*θ*) is the prior distribution of the model parameter. We assume the errors to be independent and identically distributed according to a normal distribution with an unknown standard deviation *γ* depending on the type of data. Therefore, we have the relation

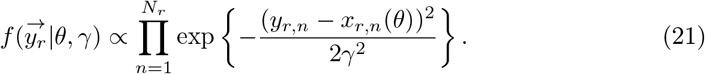

Actually, 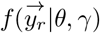 is the likelihood of the errors given the parameter *θ* and the unknown standard deviation *γ*.

Therefore, we include *γ* in the set of parameters to be estimated, meaning we seek to estimate 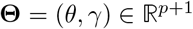. Then, we have the relations

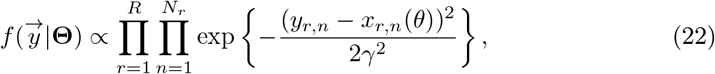

The algorithm seeks to find the parameters that maximize the probability 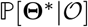. It uses the ratio,

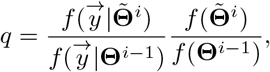

where **Θ**^*i*-1^ is the value of the parameter at the (*i* –1)-th step and 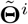 is the candidate for the *i*-th step.

In the special case where the prior distribution is chosen to be uniform, the second fraction vanishes.

Initial values for the model parameters **Θ**^0^ can be chosen using a linear programming method minimizing minus the log-likelihood function or using biological presumption. We use an Adaptive Scaling Metropolis algorithm ([63], [64]).

#### Algorithm 1: Algorithm 1

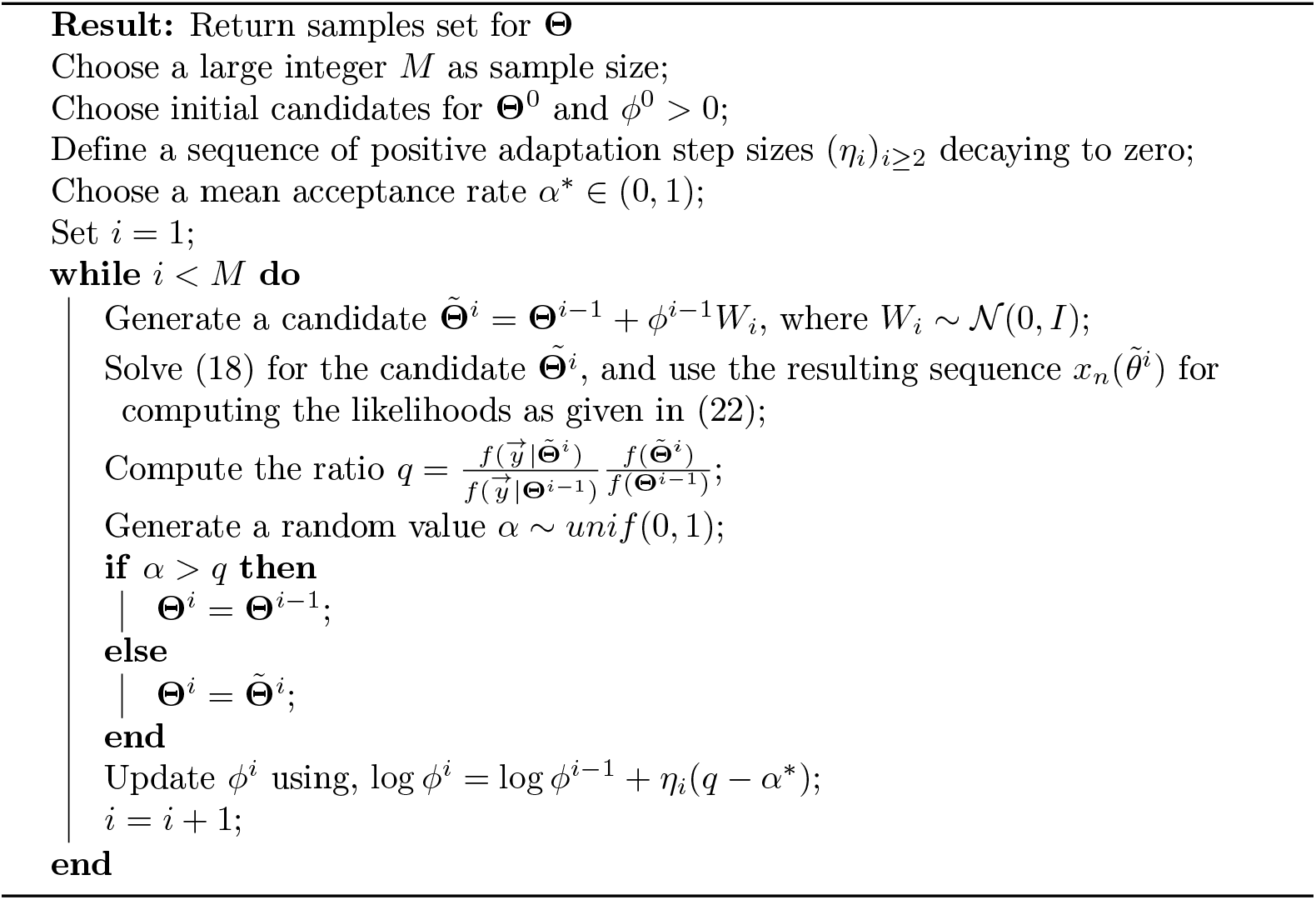

All models and simulations were implemented in MATLAB (vs.2020a, MathWorks Inc.).

#### 5.3.1 Statistical procedures

We perform a t-test in order to compare means of the different replicates in figure 6. Before performing the t-test, the assumptions of normality of the data and homogeneity of the variances were verified using the Shapiro-Wilk and Bartlett’s tests, respectively.

#### 5.3.2 Data availability

Scripts used for the models in this work can be accessed from https://github.com/IsalineLucille22/Liquid-models.git (link will be made public upon acceptance).

## Acknowledgments

The authors thank Nicolas Carraro for his help in part of the competition experiments and Noushin Hadadi for initial coding of a co-culture growth model. This work was supported by the Swiss National Science Foundation (Sinergia program, grant CRSII5 189919/1), SystemsX.ch grant 2013/158 (Design and Systems Biology of Functional Microbial Landscapes “MicroScapesX”), and by the National Centre in Competence Research in Microbiomes.

