## Supplementary information for "Regulated bacterial interaction networks: A mathematical framework to describe competitive growth under inclusion of metabolite cross-feeding"

#### Contents

**1 Supplementary figure 1: Inferred maximum growth rates of *P. veronii* on D-mannitol and *P. putida* on putrescine 2**

**2 Supplementary figure 2. Simulated biomass growth and waste formation in co-culture of *P. putida* and *P. veronii* with two independent substrates D-mannitol and putrescine. 3**

**3 Appendix S1: Logistic model for mono and co-culture growth 4**

**4 Appendix S2: Monod model for mono and co-culture growth 6**
4.1 Elimination of the intermediate complex $P$ . . . . . . . . 7
4.2 Monod model for coculture growth . . . . . . . . 8

**5 Appendix S3: Vanishing waste concentration under permanent cross-feeding 9**

**6 Appendix S4: Generalized consumer-resource model 12**

**7 Appendix S5: RBN mass action kinetics 15**

### 1 Supplementary figure 1: Inferred maximum growth rates of *P. veronii* on D-mannitol and *P. putida* on putrescine

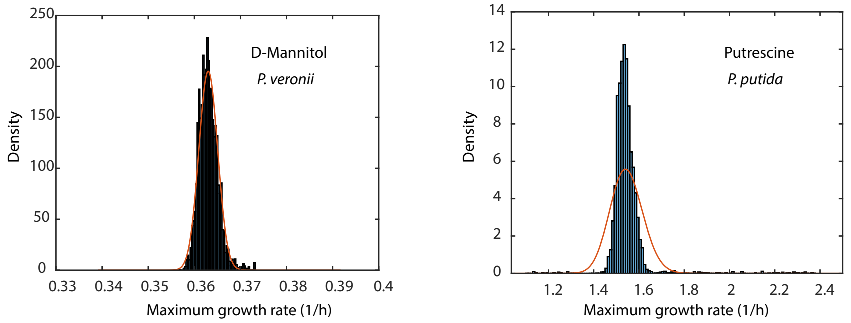


**Figure 1. Maximum growth rates of *P. veronii* and *P. putida* mono-cultures on 10 mM D-mannitol or 6.7 mM putrescine, respectively**. Plots show histograms of maximum growth rates inferred from Metropolis-Hasting fitting with Markov Chain Monte Carlo approach as described in the Materials and Methods section.

.

#### 2 Supplementary figure 2. Simulated biomass growth and waste formation in co-culture of *P. putida* and *P. veronii* with two independent substrates D-mannitol and putrescine.

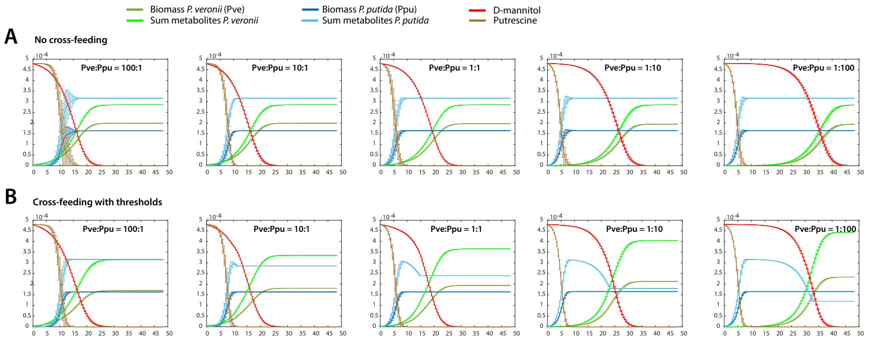


**Figure 2. Simulated biomass growth and waste formation in co-culture of *P. putida* and *P. veronii* in conditions of substrate indifference.** Plots show simulated biomass growth in 8 replicates of *P. putida* (blue) or *P. veronii* (green) in co-culture on a mixture of D-mannitol (red) and putrescine (brown), and predicted waste concentrations (light green and blue), for five different cell starting ratios (100:1, 10:1, 1:1, 1:10 and 1:100, as indicated), and $1 \times 10^6$ cells per ml at start. Simulations in (A) assume no cross-feeding in Monod model, and in (B) with cross-feeding using the discontinuous threshold function).

### 3 Appendix S1: Logistic model for mono and co-culture growth

The logistic model assumes that growth can be described as a reaction involving a species $S$ that enters in contact with a resource $R$, which it transforms to new cell biomass, and leading to cell division (duplication). Note that we use the term *species* for simplicity here, while acknowledging that the term is taxonomically poorly defined for the prokaryotic domains. This process is described by the reaction

$$S{+}R \xrightarrow{\kappa_s} 2S. \tag{1}$$

During cellular metabolism and biosynthesis, substances other than cell biomass are produced, which can leak outside the cell and which we (collectively) denote by $W$. Leaking substances may consist of regular metabolites in temporarily overflow (which the cell may take up again at a later stage), or specifically excreted compounds with a biological distinct function (e.g., signalling molecules or toxins), or wastes.
At this point, we assume that such product is a waste, which cannot be further used for growth by the same species.

$$S{+}R \xrightarrow{\kappa_w} S{+}W, \tag{2}$$

In these reactions, the kinetic rates $\kappa_s,\ \kappa_w \in \mathbb{R}^+$ refer to reaction rates; whereas $X = [S]$, $C = [R]$, $F = [W]$ are the mass concentrations of the species' biomass, of the resource and of the waste, respectively.
By using mass action law, the system of ordinary differential equations corresponding to these reactions for a mono-culture is

$$\begin{aligned} \frac{dX(t)}{dt} &= \kappa_s X(t)C(t), \\ \frac{dC(t)}{dt} &= -(\kappa_s + \kappa_w)X(t)C(t), \\ \frac{dF(t)}{dt} &= \kappa_w X(t)C(t). \end{aligned} \tag{3}$$

At any time $t$, the sum of all components $\frac{dC(t)}{dt} + \frac{dF(t)}{dt} + \frac{dX(t)}{dt} = 0$, which leads to $C(t) + F(t) + X(t) = \text{constant}$. From $\frac{dF(t)}{dt} = \frac{\kappa_w}{\kappa_s}\frac{dX(t)}{dt}$ and $\frac{dC(t)}{dt} = -\frac{\kappa_s+\kappa_w}{\kappa_s}\frac{dX(t)}{dt}$, one deduces that $F(t) = \frac{\kappa_2}{\kappa_s}X(t) + w_0$ and $C(t) = -\frac{\kappa_s+\kappa_w}{\kappa_s}X(t) + c_0$, where $w_0$ and $c_0$ are constants to be determined. The initial states of the system are $F(0) = 0$, $C(0) = C_0 > 0$ and $X(0) = X^0 > 0$ therefore, $w_0 = -\frac{\kappa_w}{\kappa_s}X^0$ and $c_0 = C_0 + \frac{\kappa_s+\kappa_w}{\kappa_s}X^0$. Because the total mass present in the system at any time $t$ is constant and equals $C(t) + F(t) + X(t) = C_0 + X^0$, we can keep a single equation to describe the system. Hence, the development of the biomass of species $S$ follows the logistic o.d.e.

$$\frac{dX(t)}{dt} = X(t)\kappa_s X_m \left(1 - \frac{X(t)}{\rho X_m}\right), \tag{4}$$

where $X_m = c_0 = C_0 + \frac{1}{\rho}X^0$ and $\rho = \frac{\kappa_s}{\kappa_s+\kappa_w}$.
We can approximate $X_m \approx C_0$ because the initial cell concentrations are negligibly low compared to the available resource concentration (in our experiments: 5 mM of succinate). We can now set the product of $\kappa_s X_m \approx \kappa_s C_0$ as $\mu_{max}$, a constant that can be interpreted as the maximum growth rate of species S in the habitat defined by the experiment. $\rho$ is a constant that represents the yield of substrate conversion into biomass for species S. The solution to (4) is then given by the standard logistic growth curve

$$X(t) = \frac{\rho X_m X^0 e^{\mu_{max}t}}{\rho X_m - X^0 + X^0 e^{\mu_{max}t}}. \tag{5}$$

##### Effect of initial concentrations for logistic coculture growth model

We can now expand the system to describe co-cultures with two different species using the same resource through the following CRN, where neither excretion products are consumed

$$\begin{aligned} S_1 + R &\xrightarrow{{\kappa_s}^1} 2S_1,\ S_1 + R \xrightarrow{{\kappa_w}^1} S_1 + W_1, \\ S_2 + R &\xrightarrow{{\kappa_s}^2} 2S_2,\ S_2 + R \xrightarrow{{\kappa_w}^2} S_2 + W_2, \end{aligned} \tag{6}$$

With the mass action law, this leads to the following set of o.d.e.s:

$$\begin{aligned} \frac{dX_1(t)}{dt} &= \kappa_s^1 X_1(t)C(t), \\ \frac{dX_2(t)}{dt} &= \kappa_s^2 X_2(t)C(t), \\ \frac{dW_1(t)}{dt} &= \kappa_w^1 X_1(t)C(t), \\ \frac{dW_2(t)}{dt} &= \kappa_w^2 X_2(t)C(t), \\ \frac{dC(t)}{dt} &= -(\kappa_s^1 + \kappa_w^1)X_1(t)C(t) - ((\kappa_s^2 + \kappa_w^2))X_2(t)C(t). \end{aligned} \tag{7}$$

We have $\frac{dW_i(t)}{dt} = \frac{\kappa_w^i}{\kappa_s^i}\frac{dX_i(t)}{dt}$ for $i = 1, 2$, therefore $W_i(t) = \frac{\kappa_w^i}{\kappa_s^i} S_i(t) + cst$. The yield for each species is defined by $\rho_i = \frac{\kappa_s^i}{\kappa_s^i + \kappa_w^i}$.
Then, if cell biomass concentrations of both species at start are negligibly low compared to the available resource, we can simplify by using the conservation relation $\frac{1}{\rho_1}X_1(t) + \frac{1}{\rho_2}X_2(t) + C(t) = \frac{1}{\rho_1}X_1^0 + \frac{1}{\rho_2}X_2^0 + C_0 \approx X_m$ from which we obtain

$$\begin{aligned} \frac{dX_1(t)}{dt} &= X_1(t)\kappa_s^1 X_m \left(1 - \frac{\frac{1}{\rho_1}X_1(t) + \frac{1}{\rho_2}X_2(t)}{X_m}\right), \\ \frac{dX_2(t)}{dt} &= X_2(t)\kappa_s^2 X_m \left(1 - \frac{\frac{1}{\rho_1}X_1(t) + \frac{1}{\rho_2}X_2(t)}{X_m}\right). \end{aligned} \tag{8}$$

By setting $\mu_{max}^i = \kappa_s^i X_m$ we obtain the system (9) of a logistic model for two species with a single unique growth resource:

$$\begin{aligned} \frac{dX_1(t)}{dt} &= X_1(t)\mu_{max}^1 \left(1 - \frac{\frac{1}{\rho_1}X_1(t) + \frac{1}{\rho_2}X_2(t)}{X_m}\right), \\ \frac{dX_2(t)}{dt} &= X_2(t)\mu_{max}^2 \left(1 - \frac{\frac{1}{\rho_1}X_1(t) + \frac{1}{\rho_2}X_2(t)}{X_m}\right). \end{aligned} \tag{9}$$

System (9) has an explicit mathematical solution. First, notice that both solutions $X_i(t)$, $i = 1, 2$ are non-decreasing and bounded, and thus converge to positive equilibria $X_i^* > 0$ when $X_i^* > 0$. Next, observe that

$$\frac{dC(t)}{dt} \leq -(\kappa_s^1 X_1^* + \kappa_s^2 X_2^*)C(t),$$

in which case the Gronwall Lemma can be used to define the upper bound

$$C(t) \leq C(0)\exp(-\beta t),$$

where $\beta = (\kappa_s^1 X_1^* + \kappa_s^2 X_2^*) > 0$. The concentration of substrate $C(t)$ decreases thus exponentially fast toward 0. The set of positive equilibria $X_i^*$ is then given by the

segment $I = \{(X_1^*, X_2^*);\ X_1^*/\rho_1 + X_2^*/\rho_2 = X_1(0)/\rho_1 + X_2(0)/\rho_2;\ X_i^* > 0,\ i = 1, 2\}$. Figure 4 of the main text shows the dependence of the limiting equilibrium from $I$ as a function of the initial biomass concentration ratio $X_1(0)/X_2(0)$.
Finally,

$$\mu^2_{max}\frac{dX_1(t)}{X_1(t)dt} = \mu^1_{max}\frac{dX_2(t)}{X_2(t)dt},$$

thus

$$\mu^2_{max}\frac{d\ln X_1(t)}{dt} = \mu^1_{max}\frac{d\ln X_2(t)}{dt}. \tag{10}$$

Solving this last equation leads to

$$X_1(t) = \frac{X_1(0)}{X_2(0)^{\frac{\mu^1_{max}}{\mu^2_{max}}}} X_2(t)^{\frac{\mu^1_{max}}{\mu^2_{max}}}.$$

We can find the steady-state $X_2^*$ as

$$x + \frac{\rho_2}{\rho_1}\frac{X_1(0)}{X_2(0)X_2(0)^{\frac{\mu^1_{max}-\mu^2_{max}}{\mu^2_{max}}}} x^{\frac{\mu^1_{max}}{\mu^2_{max}}} - \rho_2 X_m = 0. \tag{11}$$

Similarly, the steady-state of $X_1^*$ equals

$$x + \frac{\rho_1}{\rho_2}\frac{X_2(0)}{X_1(0)X_1(0)^{\frac{\mu^2_{max}-\mu^1_{max}}{\mu^1_{max}}}} x^{\frac{\mu^2_{max}}{\mu^1_{max}}} - \rho_1 X_m = 0. \tag{12}$$

Therefore, the steady-states depend on the initial ratio between the two species, and also on their initial absolute abundances, since $X_1(0)^{\frac{\mu^2_{max}-\mu^1_{max}}{\mu^1_{max}}}$ and $X_2(0)^{\frac{\mu^1_{max}-\mu^2_{max}}{\mu^2_{max}}}$. Figure 4, main text, represents the convergence to steady-state of a two species system which grow according to the chemical reactions. The different colors correspond to varying starting ratios of *P. veronii* and *P. putida* biomass. In this special case, the set of steady-states is defined by a line.

### 4 Appendix S2: Monod model for mono and co-culture growth

In contrast to the logistic model, the Monod model would assume that a bacterial species $S$ uses a unique resource $R$ to form an intermediate complex $P$, which then gives rise both to a new cell and to waste product $W$. In this case, equations (1) and (2) are rewritten as

$$\begin{aligned} S{+}R &\xrightarrow{\kappa_1} P \xrightarrow{\kappa_2} 2S, \\ P &\xrightarrow{\kappa_3} S{+}W. \end{aligned} \tag{13}$$

Let $X = [S],\ Y = [P],\ C = [R]$ and $F = [W]$ be the mass concentrations of the species $S$, the complex $P$, the resource $R$ and the waste $W$, respectively. Let $X'$ denote the mass concentration of free $S$ molecules that are not bound to $R$ molecules in complex $P$. Then the total bacterial biomass at time $t$ equals $X(t) = X'(t) + Y(t)$. From these reaction equations, we derive the following mass-action o.d.e.s

$$\begin{aligned}\frac{dX'(t)}{dt} &= -\kappa_1 X'(t)C(t) + (2\kappa_2+\kappa_3)Y(t),\\ \frac{dY(t)}{dt} &= \kappa_1 X'(t)C(t) - (\kappa_2+\kappa_3)Y(t),\\ \frac{dC(t)}{dt} &= -\kappa_1 X'(t)C(t)\\ \frac{dF(t)}{dt} &= \kappa_3 Y(t).\end{aligned} \tag{14}$$

To ensure consistency of units and denoting time in hours $[h]$, this would mean that $\kappa_2$ and $\kappa_3$ are given in $[h]^{-1}$, but that the unit of $\kappa_1$ is $([h][g]/[ml])^{-1}$.

#### 4.1 Elimination of the intermediate complex $P$

Using a quasi-steady state approximation based on $\frac{1}{\kappa_1}\frac{dY(t)}{dt} = 0$, one obtains the simpler system

$$\begin{aligned}0 &= (X(t)-Y(t))C(t) - \frac{(\kappa_2+\kappa_3)}{\kappa_1}Y(t),\\ \frac{dX(t)}{dt} &= \kappa_2 Y(t),\\ \frac{dC(t)}{dt} &= -\kappa_1(X(t)-Y(t))C(t),\\ \frac{dF(t)}{dt} &= \kappa_3 Y(t),\end{aligned} \tag{15}$$

which leads to a system with saturable consumption/degradation kinetics

$$\begin{aligned}Y(t) &= \frac{X(t)C(t)}{C(t)+\frac{\kappa_1+\kappa_2}{\kappa_1}},\\ \frac{dX(t)}{dt} &= \kappa_2\frac{X(t)C(t)}{C(t)+\frac{\kappa_2+\kappa_3}{\kappa_1}}.\\ \frac{dC(t)}{dt} &= -(\kappa_2+\kappa_3)\frac{X(t)C(t)}{C(t)+\frac{\kappa_2+\kappa_3}{\kappa_1}},\\ \frac{dF(t)}{dt} &= \kappa_3\frac{X(t)C(t)}{C(t)+\frac{\kappa_2+\kappa_3}{\kappa_1}}.\end{aligned} \tag{16}$$

As for the logistic model, $C(t)+F(t)+X(t) =$ constant and $\frac{dF(t)}{dt} = \frac{\kappa_3}{\kappa_2}\frac{dX(t)}{dt}$, $\frac{dC(t)}{dt} = -\frac{\kappa_2+\kappa_3}{\kappa_2}\frac{dX(t)}{dt}$, thus $F(t) = \frac{\kappa_3}{\kappa_2}X(t)+w_0$ and $C(t) = -\frac{\kappa_2+\kappa_3}{\kappa_{i2}}X(t)+c_0$ where $w_0$, $c_0$ are constants to be determined. By using the initial conditions $F(0)=0$, $C(0)=C_0>0$ and $X(0)=X^0>0$, this leads to $w_0 = -\frac{\kappa_3}{\kappa_2}X^0$ and $c_0 = C_0+\frac{\kappa_2+\kappa_3}{\kappa_2}X^0$. The total mass present in the system at any time $t$ is constant and equals $C(t)+F(t)+X(t) = C_0+X^0$. The development of the biomass of the species $S$ then follows:

$$\frac{dX(t)}{dt} = X(t)\kappa_2\frac{C_0+\frac{1}{\rho}X(0)-\frac{1}{\rho}X(t)}{C_0+\frac{1}{\rho}X(0)-\frac{1}{\rho}X(t)+\frac{\kappa_2+\kappa_3}{\kappa_1}}, \tag{17}$$

If we name $\rho = \frac{\kappa_2}{\kappa_2+\kappa_3}$, $X_m = c_0 = C_0+\frac{1}{\rho}X^0$, $\mu_{max} = \kappa_2$ and $K_S = \frac{\kappa_2+\kappa_3}{\kappa_1}$ this leads to the Monod-model of substrate to biomass growth (18).

$$\frac{dX(t)}{dt} = \mu_{max}X(t)\frac{X_m-\frac{1}{\rho}X(t)}{X_m-\frac{1}{\rho}X(t)+K_S}, \tag{18}$$

where $K_S$ is the resource concentration at which the growth rate of species $S$ is half maximal.

#### 4.2 Monod model for coculture growth

In the case of two bacterial species (denoted by $S_1$ and $S_2$) and a single common resource ($R$) we obtain the CRNs (4) of the main text

$$\begin{aligned} S_1 + R &\xrightarrow{\kappa_{1_1}} P_1 \xrightarrow{\kappa_{1_2}} 2S_1,\ P_1 \xrightarrow{\kappa_{1_3}} S_1 + W_1, \\ S_2 + R &\xrightarrow{\kappa_{2_1}} P_2 \xrightarrow{\kappa_{2_2}} 2S_2,\ P_2 \xrightarrow{\kappa_{2_3}} S_2 + W_2. \end{aligned} \tag{19}$$

Proceeding the same way as for the system with one species and using the quasi-steady state approximation ($\frac{1}{\kappa_{i_1}}\frac{dY_i(t)}{dt} = 0$) we obtain the o.d.e.s:

$$\begin{aligned} Y_1(t) &= \frac{X_1(t)C(t)}{C(t) + \frac{\kappa_{1_2}+\kappa_{1_3}}{\kappa_{1_1}}},\ Y_2(t) = \frac{X_2(t)C(t)}{C(t) + \frac{\kappa_{2_2}+\kappa_{2_3}}{\kappa_{2_1}}}, \\ \frac{dX_1(t)}{dt} &= \kappa_{1_2}\frac{X_1(t)C(t)}{C(t) + \frac{\kappa_{1_2}+\kappa_{1_3}}{\kappa_{1_1}}},\ \frac{dX_2(t)}{dt} = \kappa_{2_2}\frac{X_2(t)C(t)}{C(t) + \frac{\kappa_{2_2}+\kappa_{2_3}}{\kappa_{2_1}}}, \\ \frac{dC(t)}{dt} &= -(\kappa_{1_2}+\kappa_{1_3})\frac{X_1(t)C(t)}{C(t) + \frac{\kappa_{1_2}+\kappa_{1_3}}{\kappa_{1_1}}} - (\kappa_{2_2}+\kappa_{2_3})\frac{X_2(t)C(t)}{C(t) + \frac{\kappa_{2_2}+\kappa_{2_3}}{\kappa_{2_1}}}, \\ \frac{dF(t)}{dt} &= \kappa_{1_3}\frac{X_1(t)C(t)}{C(t) + \frac{\kappa_{1_2}+\kappa_{1_3}}{\kappa_{1_1}}} + \kappa_{2_3}\frac{X_2(t)C(t)}{C(t) + \frac{\kappa_{2_2}+\kappa_{2_3}}{\kappa_{2_1}}}. \end{aligned} \tag{20}$$

where $X_i(t) = \tilde{X}_i(t) + Y_i(t),\ \tilde{X}_i = [S_i],\ Y_i = [P_i]$ and $C = [R]$. This kind of co-culture growth model with saturable degradation/consumption kinetics appears, e.g., in [1].

As before, we can make some simplifications because the variations of species, waste and resource concentrations are combinations of $Y_1(t)$ and $Y_2(t)$. Since $\frac{dC(t)}{dt} = -\frac{\kappa_{1_2}+\kappa_{1_3}}{\kappa_{1_2}}\frac{dX_1(t)}{dt} - \frac{\kappa_{2_2}+\kappa_{2_3}}{\kappa_{2_2}}\frac{dX_2(t)}{dt}$ this leads to $C(t) = -\frac{\kappa_{1_2}+\kappa_{1_3}}{\kappa_{1_2}}X_1(t) - \frac{\kappa_{2_2}+\kappa_{2_3}}{\kappa_{2_2}}X_2(t) + c_0$. Under the initial conditions $C(0) = C_0$ and $X_i(0) = X_i^0$, $c_0 = C_0 + \frac{\kappa_{1_2}+\kappa_{1_3}}{\kappa_{1_2}}X_1^0 + \frac{\kappa_{2_2}+\kappa_{2_3}}{\kappa_{2_2}}X_2^0$. By substituting $\mu^i_{max} = \kappa_{i_2}$, $\rho_1 = \frac{\kappa_{1_2}}{\kappa_{1_2}+\kappa_{1_3}}$ and $\rho_2 = \frac{\kappa_{2_2}}{\kappa_{2_2}+\kappa_{2_3}}$, we can simplify $c_0 = C_0 + \frac{1}{\rho_1}X_1^0 + \frac{1}{\rho_2}X_2^0$ that we call $X_m$, and further writing $K_S^{(i)} = \frac{\kappa_{i_2}+\kappa_{i_3}}{\kappa_{i_1}}$, we finally obtain

$$\begin{aligned} \frac{dX_1(t)}{dt} &= X_1(t)\mu^1_{max}\left(\frac{X_m - \frac{1}{\rho_1}X_1(t) - \frac{1}{\rho_2}X_2(t)}{X_m - \frac{1}{\rho_1}X_1(t) - \frac{1}{\rho_2}X_2(t) + K_S^{(1)}}\right), \\ \frac{dX_2(t)}{dt} &= X_2(t)\mu^2_{max}\left(\frac{X_m - \frac{1}{\rho_1}X_1(t) - \frac{1}{\rho_2}X_2(t)}{X_m - \frac{1}{\rho_1}X_1(t) - \frac{1}{\rho_2}X_2(t) + K_S^{(2)}}\right). \end{aligned} \tag{21}$$

In the special condition where $K_S^{(1)} = K_S^{(2)}$, we obtain again the relation (10) and then the steady-states for the species $S_1$ and $S_2$ can be found by solving equations (12) and (11), respectively.

### 5 Appendix S3: Vanishing waste concentration under permanent cross-feeding

Let $I(t) = (F_1(t), F_2(t), C(t))$ be the line vector giving the concentrations of species $W_1$, $W_2$ and $R$ at time $t > 0$. The mass action o.d.e. associated to the CRNs (7,8) of the main text is given by the sum of the o.d.e. associated to the pairwise Lotka-Volterra system (see Figure 1 A), main text), and of catalytic conversion reactions (see Figure 1 B, main text). These conversion reactions can be described by a Laplace matrix $L_{cat}$ where the entries are related to species $W_1$, $W_2$ and $R$, which are functions of the species $S_1$ and $S_2$ biomass concentrations $X_1$ and $X_2$:

$$L_{cat}(X_1, X_2) = \begin{bmatrix} -\tilde{\kappa}_{12}X_2 & \tilde{\kappa}_{12}X_2 & 0 \\ \tilde{\kappa}_{21}X_1 & -\tilde{\kappa}_{21}X_1 & 0 \\ \tilde{\kappa}_1 X_1 & \tilde{\kappa}_2 X_2 & -\tilde{\kappa}_1 X_1 - \tilde{\kappa}_2 X_2 \end{bmatrix}.$$

One can then define the matrix $Q = L_{cat} - D$, where $D$ is the diagonal matrix

$$D(X_1, X_2) = \begin{bmatrix} \kappa_{12}X_2 & 0 & 0 \\ 0 & \kappa_{21}X_1 & 0 \\ 0 & & \kappa_1 X_1 + \kappa_2 X_2 \end{bmatrix},$$

which accounts for the consumption of species $W_1$, $W_2$ and $R$ from $S_1$ and $S_2$. The mass action o.d.e. describing the time evolution of $I(t)$ is then given by

$$\frac{dI(t)}{dt} = I(t)Q(X_1(t), X_2(t)). \tag{22}$$

The development of species $S_1$ and $S_2$ biomass concentrations over time is then presented by

$$\begin{aligned} \frac{dX_1(t)}{dt} &= X_1(t)(\kappa_1 C(t) + \kappa_{21}F_2(t)), \\ \frac{dX_2(t)}{dt} &= X_2(t)(\kappa_2 C(t) + \kappa_{12}F_1(t)). \end{aligned} \tag{23}$$

The mass action o.d.e. associated with the CRNs (7,8) of the main text is given by the system of o.d.e. (22,23). This system is conservative so that the biomass $X_i(t)$, $i = 1, 2$ remains bounded. The derivative of these functions being non-negative it follows that both biomass concentrations converge toward positive limiting values $X_i(\infty) > 0$.

**Proposition 1.** The solutions $X_i(t)$ with $X_i(0) > 0$, $i = 1, 2$ of the mass action o.d.e. (22,23) are non-decreasing and bounded, and converge toward positive limits $X_i(\infty) > 0$. There exists a positive constant $\lambda > 0$ such that $C(t) \leq C(0)\exp(-\lambda t)$. Consider the linear o.d.e. (22), where the concentration functions $X_1(t)$, $X_2(t)$, $F_1(t)$, $F_2(t)$ and $C(t)$ solve the mass-action o.d.e. (22,23). Then the origin $I = 0$ is globally asymptotically stable for the positive orthant. The manifold of states $(X_1, X_2, F_1 = 0, F_2 = 0, C = 0)$ with $X_1 + X_2 = C(0) + X_1(0) + X_2(0)$ is an equilibrium manifold.

Proposition 1, which is proven in Appendix S3, shows that, starting from any initial condition for the various concentrations with positive $X_i(0) > 0$, the waste concentrations $F_i(t)$ and $C(t)$ will ultimately converge toward 0, so that the resulting consumer steady-state concentrations $X_i(t)$ will converge toward some $X_i^*$ with $X_1^* + X_2^* = X_1(0) + X_2(0) + C(0)$. Hence, the cross-feeding mechanism of the generalized consumer-resource model (see [2, 3])) leads to vanishing waste steady-state concentrations.

The linear equation obtained by setting $dX_i/dt = 0$ further implies that the critical points with $X_i > 0$ are such that $C = F_1 = F_2 = 0$. This small argument might lead to a misunderstanding about the true content of Proposition 1. The convergence of all orbits towards equilibrium like $(X_1^*, X_2^*, 0, 0, 0)$ follows from the structure of the matrix $Q = L_{cat} - D$: $L_{cat}$ is a Laplace matrix which preserves mass, that is, describes how metabolite molecules are transformed into another metabolite, while $D$ describes consumption of metabolite molecules by consumers, and thus induces total metabolite mass decrease. We consider a more general model by adding arbitrary nonlinear terms $f_i(Z(t))$, $i = 1, 2$, to (23), where $Z(t) = (X_1(t), X_2(t), C(t), F_1(t), F_2(t))$. For example, we might also consider competition between consumers, and these new functions might take the form

$$f_i(Z(t)) = -\sum_{j \neq i} C_{ij}^{comp} X_i(t) X_j(t) + r_i X_i(t),$$

for competition coefficients $C_{ij}^{comp} > 0$ and intrinsic growth rates $r_i$. Another possible model might include predation between consumers. The new version of (23) is

$$\begin{aligned} \frac{dX_1(t)}{dt} &= X_1(t)(\kappa_1 C(t) + \kappa_{21} F_2(t)) + f_1(Z(t)), \\ \frac{dX_2(t)}{dt} &= X_2(t)(\kappa_2 C(t) + \kappa_{12} F_1(t)) + f_1(Z(t)). \end{aligned} \tag{24}$$

**Proposition 2.** Assume that the o.d.e. (22 and 24) admits unique solutions such that $X_1$ and $X_2$ converge toward positive limiting values. Let $I(t) = (F_1(t), F_2(t), C(t))$ and consider the linear o.d.e. (22), where the concentration function $Z(t)$ solves the mass-action o.d.e. (22,24). Then the origin $I = 0$ is globally asymptotically stable for the positive orthant.

#### Proof of Proposition 1 and 2:

We begin with the proof of Proposition 1: A simple analysis of the mass-action o.d.e. associated with the CRN reveals that the equilibria $(X_1, X_2, C, F_1, F_2)$ are such that $C = 0$. We thus study the solutions of (23).One can check that manifolds of equilibria exist where either $X_1 = 0$ or $X_2 = 0$, but our equations lead that $dX_i(t)/dt \geq 0$, so that imposing initial conditions with $X_i(0) > 0$, $i = 1, 2$, the orbits of the o.d.e. will never reach theses invariant manifolds and, using the fact that they are bounded and non-decreasing (see, e.g. [4]), converge toward positive equilibria $X_i(\infty) > 0$.
We then show that $C(t)$ converges exponentially fast toward 0 as $t \to \infty$. We have seen that $\lim_{t\to\infty} X_i(t) = X_i(\infty) > X_i(0)$, and that $X_i(t)$ is non-decreasing, $i = 1, 2$, so that

$$\begin{aligned} \frac{dC(t)}{dt} &= -((\kappa_1 + \kappa_1) X_1(t) + (\kappa_2 + \kappa_2) X_2(t)) C(t) \\ &\leq -(((\kappa_1 + \kappa_1) X_1(0) + (\kappa_2 + \kappa_2) X_2(t)) C(t) = -\lambda C(t), \end{aligned}$$

with $C(t) > 0$, $\forall t \geq 0$, where $\lambda = (\kappa_1 + \kappa_1) X_1(0) + (\kappa_2 + \kappa_2) X_2(0) > 0$. Gronwall's inequality then yields the upper bound

$$C(t) \leq C(0) \exp(-\lambda t), \; t \geq 0.$$

We use these estimates to prove that the origin is globally asymptotically stable for the o.d.e. (22) on the positive orthant.
The following arguments finish the proof of Proposition 1 and prove Proposition 2. The CRN given in (7,8) of the main text contains a subsystem involving catalytic conversion, see Fig. 1 B, main text, whose dynamics is well described by a Laplace matrix $L_{cat}$

where the entries are related to species $W_1$, $W_2$ and $R$, which depends on the $S_1$ and $S_2$ concentrations $X_1$ and $X_2$

$$L_{cat}(X_1, X_2) = \begin{bmatrix} -\tilde{\kappa}_{12}X_2 & \tilde{\kappa}_{12}X_2 & 0 \\ \tilde{\kappa}_{21}X_1 & -\tilde{\kappa}_{21}X_1 & 0 \\ \tilde{\kappa}_1 X_1 & \tilde{\kappa}_2 X_2 & -\tilde{\kappa}_1 X_1 - \tilde{\kappa}_2 X_2 \end{bmatrix}.$$

As seen previously, both $X_i(t)$ converge toward positive limits $X_i(\infty)$, so that one can consider the limiting Laplace matrix

$$L_{cat,\infty} = \begin{bmatrix} -\tilde{\kappa}_{12}X_2(\infty) & \tilde{\kappa}_{12}X_2(\infty) & 0 \\ \tilde{\kappa}_{21}X_1(\infty) & -\tilde{\kappa}_{21}X_1(\infty) & 0 \\ \tilde{\kappa}_1 X_1(\infty) & \tilde{\kappa}_2 X_2(\infty) & -\tilde{\kappa}_1 X_1 - \tilde{\kappa}_2 X_2(\infty) \end{bmatrix},$$

and, similarly, one arrives at the limiting diagonal matrix

$$D_\infty = \begin{bmatrix} \kappa_{12}X_2(\infty) & 0 & 0 \\ 0 & \kappa_{21}X_1(\infty) & 0 \\ 0 & & \kappa_1 X_1(\infty) + \kappa_2 X_2(\infty) \end{bmatrix},$$

Let $I(t) = (I_1(t), I_2(t), I_3(t)) = (F_1(t), F_2(t), C(t))$ be the line vector giving their concentrations at time $t > 0$.The mass action kinetics leads to

$$\frac{dI(t)}{dt} = I(t)Q, \tag{25}$$

where $Q = L_{cat}(X_1(t), X_2(t)) - D$.
Using the convergence of $X_i(t)$, this last o.d.e. is close to the linear autonomous o.d.e.

$$\frac{dI(t)}{dt} = I(t)Q_\infty, \tag{26}$$

where $Q_\infty = L_{cat,\infty} - D_\infty$.
The structure of the o.d.e. (25) suggests that its asymptotic behaviour is close to that of the autonomous one. First notice that

$$\frac{dI(t)}{dt} = I(t)Q_\infty + g(t, I(t)),$$

where

$$g(t, y) = y(Q(t) - Q_\infty).$$

The entries of the matrix $Q(t) - Q_\infty$ are products involving the various kinetic constants or sums of them, that are bounded by some positive constant $K^0 > 0$, so that

$$||g(t, y)|| \leq 3K^0 \sup_k |y_k| \max\{|X_1(t) - X_1(\infty)|, |X_2(t) - X_2(\infty)|\}.$$

It follows that, using the convergence of the $X_i$ and the boundedness of all functions, we arrive at $||g(t, y)|| \leq \gamma(t)$, for all $y$ in a compact $B$ that depend on the initial concentrations, with $\gamma(t) \to 0$ as $t \to \infty$. Then Corolloray 3.3. of [5] shows that all solutions of (25) converge toward 0 when the origin is globally asymptotically stable for the autonomous o.d.e., as required for Proposition 1.
To show that the origin is globally asymptotically stable for the o.d.e. (26), is it sufficient to prove that all the real parts of the eigenvalues of the matrix $Q_\infty$ are negative. This is seen using the Levy-Hadamard Theorem, which states that all of the eigenvalues are contained in the union of the discs of center $(Q_\infty)_{kk}$ of radius $\sum_{m \neq k} (Q_\infty)_{km}$. A direct computation shows that this holds true.

### 6 Appendix S4: Generalized consumer-resource model

We show that the mass-action o.d.e. associated to the CRN (7,8) of the main text

$$S_1 + R \xrightarrow{\kappa_1} 2S_1, \;\; S_1 + R \quad \xrightarrow{\tilde{\kappa}_1} S_1 + W_1,$$
$$S_2 + R \xrightarrow{\kappa_2} 2S_2, \;\; S_2 + R \quad \xrightarrow{\tilde{\kappa}_2} S_2 + W_2,$$

$$S_1 + W_2 \xrightarrow{\kappa_{21}} 2S_1, \quad S_1 + W_2 \quad \xrightarrow{\tilde{\kappa}_{21}} S_1 + W_1,$$
$$S_2 + W_1 \xrightarrow{\kappa_{12}} 2S_2, \quad S_2 + W_1 \quad \xrightarrow{\tilde{\kappa}_{12}} S_2 + W_2.$$

can be recast as a generalized consumer-resource model, but involves fewer parameters. The mass action kinetics o.d.e. associated with (7,8) is given by

$$\begin{aligned}
\frac{dX_1(t)}{dt} &= \kappa_1 X_1(t)F_3(t) + \kappa_{21}X_1(t)F_2(t),\\
\frac{dX_2(t)}{dt} &= \kappa_2 X_2(t)F_3(t) + \kappa_{12}X_2(t)F_1(t),\\
\frac{dF_1(t)}{dt} &= \tilde{\kappa}_1 X_1(t)F_3(t) - (\kappa_{12} + \tilde{\kappa}_{12})X_2(t)F_1(t) + \tilde{\kappa}_{21}X_1(t)F_2(t),\\
\frac{dF_2(t)}{dt} &= \tilde{\kappa}_2 X_2(t)F_3(t) - (\kappa_{21} + \tilde{\kappa}_{21})X_1(t)F_2(t) + \tilde{\kappa}_{12}X_2(t)F_1(t),\\
\frac{dF_3(t)}{dt} &= -(\kappa_1 + \tilde{\kappa}_1)X_1(t)F_3(t) - (\kappa_2 + \tilde{\kappa_2})X_2(t)F_3(t),
\end{aligned} \tag{27}$$

where $X_i$ are the concentrations of species $S_i$, $i = 1, 2$, and where $F_k$, $k = 1, 2, 3$ are the concentrations of waste $W_k$, $k = 1, 2$ and of resource $R$.
The network structure (see Fig. 1 of the main text) suggests that the secretion matrix $P$ is of the form

$$P = \begin{bmatrix} 0 & p_{12} & 0 \\ p_{21} & 0 & 0 \\ p_{31} & p_{32} & 0 \end{bmatrix}$$

The assumed stochasticity of the secretion matrix imposes that $p_{12} = p_{21} = 1$ and $p_{31} + p_{32} = 1$, and the resulting generalized consumer-resource model becomes

$$\begin{aligned}
\frac{dN_1}{dt} &= g_1 N_1((1 - l_3)q_5 c_{13} R_3 + (1 - l_2)q_2 c_{12} R_2),\\
\frac{dN_2}{dt} &= g_2 N_2((1 - l_3)q_3 c_{23} R_3 + (1 - l_1)q_1 c_{21} R_1),\\
\frac{dR_1}{dt} &= -c_{21} R_1 N_2 + \Lambda_1,\\
\frac{dR_2}{dt} &= -c_{12} R_2 N_1 + \Lambda_2,\\
\frac{dR_3}{dt} &= -c_{13} R_3 N_1 - c_{23} R_3 N_2,
\end{aligned} \tag{28}$$

where

$$\Lambda_1 = l_2 \frac{q_2}{q_1} c_{12} R_2 N_1 + l_3 p_{31} \frac{q_3}{q_1} c_{13} R_3 N_1,$$

$$\Lambda_2 = l_1 \frac{q_1}{q_2} c_{21} R_1 N_2 + l_3 p_{32} \frac{q_3}{q_2} c_{23} R_3 N_2,$$

aand $\Lambda_3 = 0$. One then compares the two o.d.e.s to arrive at the two sets of equations which correspond to Fig 1 A and B of the main text:

$$\kappa_1 = g_1(1 - l_3)q_3 c_{13}, \quad \kappa_{21} = g_1(1 - l_2)q_2 c_{12},$$

$$\kappa_2 = g_2(1-l_3)q_3c_{23}, \quad \kappa_{12} = g_2(1-l_1)q_1c_{21},$$

and

$$\tilde{\kappa}_1 = l_3p_{31}\frac{q_3}{q_1}c_{13}, \quad \tilde{\kappa}_{21} = l_2\frac{q_2}{q_1}c_{12},$$

$$\tilde{\kappa}_2 = l_2p_{32}\frac{q_3}{q_2}c_{23}, \quad \tilde{\kappa}_{12} = l_1\frac{q_1}{q_2}c_{21}.$$

Suppose we are given the parameters of (27). Then we describe the set of parameters $g_1$, $g_2$, $l_k$, $q_k$ and $c_{ik}$ that solve the above set of equations. We first observe that necessarily

$$c_{13} = \kappa_1 + \tilde{\kappa}_1, \quad c_{23} = \kappa_2 + \tilde{\kappa}_2,$$

$$c_{21} = \kappa_{12} + \tilde{\kappa}_{12}, \quad c_{12} = \kappa_{21} + \tilde{\kappa}_{21}.$$

We then focus on the second set of equations using notions from algebraic potential theory associated to the directed graph given in Fig. ?? of node set $V = \{R_1,\ R_2,\ R_3\}$ and edge set $E = \{f_1,\ f_2,\ f_3,\ f_4\}$. Set for convenience

$$y_1^{-1} = \frac{\tilde{\kappa}_1}{l_3p_{31}c_{13}}, \quad y_2^{-1} = \frac{\tilde{\kappa}_{21}}{l_2c_{12}}, \quad y_3^{-1} = \frac{\tilde{\kappa}_2}{l_3p_{32}c_{23}}, \quad y_4^{-1} = \frac{\tilde{\kappa}_{12}}{l_1c_{21}}.$$

Let $L_y = (\ln(y_1), \ln(y_2), \ln(y_3), \ln(y_4))^T$, and consider the incidence matrix

$$I = \begin{bmatrix} 1 & 1 & 0 & -1 \\ 0 & -1 & 1 & 1 \\ -1 & 0 & -1 & 0 \end{bmatrix}$$

Then the second block of equation becomes

$$L_y = I^T L_q, \tag{29}$$

where $L_q = (\ln(q_1), \ln(q_2), \ln(q_3))^T$, which corresponds to a Dirichlet problem, see [6]. The last equation has a solution when $L_y \in \mathrm{Im}(I^T)$, which is the orthogonal complement to the cycle space of the graph (Proposition 4.1 of [6]). The two cycles of the graph are $(f_4, f_2)$ and $(f_1, f_4, -f_3)$ of characteristic vectors $(0, 1, 0, 1)$ and $(1, 0, -1, 1)$. The orthogonality relation yields the two conditions

$$\frac{\tilde{\kappa}_{21}}{\tilde{\kappa}_{21} + \kappa_{21}}\frac{\tilde{\kappa}_{12}}{\tilde{\kappa}_{12} + \kappa_{12}} = l_1l_2, \tag{30}$$

and

$$\frac{\tilde{\kappa}_1}{\tilde{\kappa}_1 + \kappa_1}\frac{\tilde{\kappa}_2 + \kappa_2}{\tilde{\kappa}_2}\frac{\tilde{\kappa}_{12}}{\tilde{\kappa}_{12} + \kappa_{12}} = \frac{p_{31}}{p_{32}}l_1. \tag{31}$$

From here, one can solve explicitly (29) using the general theory as in [6], Section 9, or work directly to arrive at the solutions when both (30) and (31) are satisfied.
The parameters $l_1$, $l_2$, $q_1$ and $q_3$ can be deduced from the other. Set for convenience

$$q_{21} = \frac{\tilde{\kappa}_{21}}{\tilde{\kappa}_{21} + \kappa_{21}}, \quad q_{12} = \frac{\tilde{\kappa}_{12}}{\tilde{\kappa}_{12} + \kappa_{12}},$$

and

$$q_{31} = \frac{\tilde{\kappa}_1}{\tilde{\kappa}_1 + \kappa_1}, \quad q_{32} = \frac{\tilde{\kappa}_2}{\tilde{\kappa}_2 + \kappa_2}.$$

Hence, with these notations,

$$l_1 = \frac{q_{31}q_{12}}{q_{32}}\frac{p_{32}}{p_{31}}, \tag{32}$$

and

$$l_2 = \frac{q_{21}q_{32}}{q_{31}}\frac{p_{31}}{p_{32}}, \quad (33)$$

$$q_1 = \frac{q_2 l_2}{q_{21}}, \quad (34)$$

and

$$q_3 = \frac{q_{32}}{p_{32}l_3}q_2. \quad (35)$$

One can then focus on the first set of equations, which becomes after some computations,

$$\frac{\kappa_1}{c_{13}q_{32}} = g_1\frac{1-l_3}{l_3}\frac{q_2}{p_{32}},$$
$$\frac{\kappa_{21}}{c_{12}} = g_1(1-\frac{q_{21}q_{32}}{q_{31}}\frac{p_{31}}{1-p_{31}})q_2,$$
$$\frac{\kappa_2}{c_{23}q_{32}} = g_2\frac{1-l_3}{l_3}\frac{q_2}{p_{32}},$$
$$\frac{\kappa_{12}q_{21}}{c_{21}} = g_2(\frac{q_{21}q_{32}}{q_{31}}\frac{p_{31}}{p_{32}} - q_{21}q_{12})q_2.$$

Let A, B, C and D be the following functions of $p_{31}$ (recall that $p_{31}+p_{32}=1$)

$$A = \frac{\kappa_1}{c_{13}q_{32}}(1-p_{31}), \quad B = \frac{\kappa_{21}}{c_{12}}\frac{1}{1-\frac{q_{21}q_{32}}{q_{31}}\frac{p_{31}}{1-p_{31}}},$$

$$C = \frac{\kappa_2}{c_{23}q_{32}}(1-p_{31}), \quad D = \frac{\kappa_{12}q_{21}}{c_{21}}\frac{1}{(\frac{q_{21}q_{32}}{q_{31}}\frac{p_{31}}{p_{32}} - q_{21}q_{12})}.$$

The equation is then equivalent to

$$A = g_1(\frac{1}{l_3}-1)q_2, \quad B = g_1q_2, \quad C = g_2(\frac{1}{l_3}-1)q_2, \quad D = g_2q_2$$

Considering the ratios $A/B$ and $C/D$, one deduces the relations

$$l_3 = \frac{1}{1+\frac{A}{B}} = \frac{1}{1+\frac{C}{D}}. \quad (36)$$

One thus must check that there is some $p_{31}$ satisfying (36) When $A \neq -B$ and $C \neq -D$, this is equivalent to $\frac{A}{B} = \frac{C}{D}$. which corresponds to :

$$\frac{\kappa_1}{\kappa_1+\tilde{\kappa}_1}\frac{\kappa_2+\tilde{\kappa}_2}{\tilde{\kappa}_2}\frac{\kappa_{21}+\tilde{\kappa}_{21}}{\kappa_{21}}(1-p_{31}) - \frac{\tilde{\kappa}_{21}}{\kappa_{21}} = \frac{\kappa_2^2}{\tilde{\kappa}_2(\tilde{\kappa}_2+\kappa_2)}\frac{\kappa_{12}+\tilde{\kappa}_{12}}{\kappa_{12}}\frac{\tilde{\kappa}_1+\kappa_1}{\tilde{\kappa}_1}p_{31} - \frac{\kappa_2\tilde{\kappa}12}{\tilde{\kappa}_2\kappa_{12}}(1-p_{31}),$$

wich can be solved to arrive at

$$p_{31} = \frac{\frac{\kappa_1}{\kappa_1+\tilde{\kappa}_1}\frac{\kappa_2+\tilde{\kappa}_2}{\tilde{\kappa}_2}\frac{\kappa_{21}+\tilde{\kappa}_{21}}{\kappa_{21}} + \frac{\kappa_2\tilde{\kappa}_{12}}{\tilde{\kappa}_2\kappa_{12}}}{\frac{\kappa_1}{\kappa_1+\tilde{\kappa}_1}\frac{\kappa_2+\tilde{\kappa}_2}{\tilde{\kappa}_2}\frac{\kappa_{21}+\tilde{\kappa}_{21}}{\kappa_{21}} + \frac{\tilde{\kappa}_{21}}{\kappa_{21}} + \frac{\kappa_2^2}{\tilde{\kappa}_2(\tilde{\kappa}_2+\kappa_2)}\frac{\kappa_{12}+\tilde{\kappa}_{12}}{\kappa_{12}}\frac{\tilde{\kappa}_1+\kappa_1}{\tilde{\kappa}_1} + \frac{\kappa_2\tilde{\kappa}_{12}}{\tilde{\kappa}_2\kappa_{12}}},$$

Hence,

$$g_1 = \frac{A}{q_2(\frac{1}{l_3}-1)}, \quad g_2 = \frac{D}{q_2}.$$

One sees that 1) the constants $g_i$, $i=1,2$ are necessary and that 2) the model is not identifiable since this last system has a one dimensional solution set.

### 7 Appendix S5: RBN mass action kinetics

We consider the following simplified CRN given in (11,12) of the main text

$$
\begin{aligned}
&S_1 + R \xrightarrow{\kappa_1} 2S_1,\ S_1 + R \quad \xrightarrow{\tilde{\kappa}_1} S_1 + W_1, T \\
&S_2 + R \xrightarrow{\kappa_2} 2S_2,\ S_2 + R \qquad \xrightarrow{\tilde{\kappa}_2} S_2 + W_2,
\end{aligned} \tag{37}
$$

$$
\begin{aligned}
&S_1 + W_2 \xrightarrow{\phi(\mathrm{F}_2 - \mathrm{T}_{\mathrm{W}_1})\kappa_{21}} 2S_1, \quad S_1 + W_2 \quad \xrightarrow{\phi(\mathrm{F}_2 - \mathrm{T}_{\mathrm{W}_1})\tilde{\kappa}_{21}} S_1 + W_1, \\
&S_2 + W_1 \xrightarrow{\phi(\mathrm{F}_1 - \mathrm{T}_{\mathrm{W}_2})\kappa_{12}} 2S_2, \quad S_2 + W_1 \quad \xrightarrow{\phi(\mathrm{F}_2 - \mathrm{T}_{\mathrm{W}_2})\tilde{\kappa}_{12}} S_2 + W_2,
\end{aligned} \tag{38}
$$

of associated mass-action o.d.e.

$$
\begin{aligned}
\frac{dX_1(t)}{dt} &= \kappa_1 X_1(t)C(t) + \phi(F_2 - T_{W_1})\kappa_{21} X_1(t)F_2(t), \\
\frac{dX_2(t)}{dt} &= \kappa_2 X_2(t)C(t) + \phi(F_1 - T_{W_2})\kappa_{12} X_2(t)F_1(t), \\
\frac{dF_1(t)}{dt} &= \tilde{\kappa}_1 X_1(t)C(t) - \phi(F_1 - T_{W_2})(\kappa_{12} + \tilde{\kappa}_{12})X_2(t)F_1(t) + \phi(F_2 - T_{W_1})\tilde{\kappa}_{21} X_1(t)F_2(t), \\
\frac{dF_2(t)}{dt} &= \tilde{\kappa}_2 X_2(t)C(t) - \phi(F_2 - T_{W_1})(\kappa_{21} + \tilde{\kappa}_{21})X_1(t)F_2(t) + \phi(F_1 - T_{W_2})\tilde{\kappa}_{12} X_2(t)F_1(t), \\
\frac{dC(t)}{dt} &= -(\kappa_1 + \tilde{\kappa}_1)X_1(t)C(t) - (\kappa_2 + \tilde{\kappa_2})X_2(t)C(t).
\end{aligned} \tag{39}
$$

Assume e.g. that the activation function of the RBN is $\phi_\varepsilon(x) = x/(2\varepsilon) + 1/2$ for $|x| \leq \varepsilon$, $\phi_\varepsilon(x) = 0$ for $x \leq -\varepsilon$ and $\phi_\varepsilon(x) = 1$ when $x \geq \varepsilon$. We look for critical points $(X_1^*, X_2^*, F_1^*, F_2^*, C^*)$ of the o.d.e. (39) with $X_i^* > 0$, $i = 1, 2$. Such critical point must satisfy

$$\kappa_1 X_1^* C^* + \phi(F_2^* - T_{W_1})\kappa_{21} X_1^* F_2^* = 0.$$

The solution of mass action being non-negative when starting from non-negative initial conditions (see [4]), we get that necessarily $C^* = 0$ and that either $F_2^* = 0$ or $\phi_\varepsilon(F_2^* - T_{W_1}) = 0$, that is $F_2^* \leq T_{W_1} - \varepsilon$. Proceeding similarly for the remaining equations, one arrives at the following manifold $I$ of equilibria

$$I = \{(X_1^*, X_2^*, F_1^*, F_2^*, C^*);\ X_i^* > 0,\ F_i^* \leq T_i - \varepsilon,\ i = 1,\ 2,\ C^* = 0\}.$$
